# Towards Understanding the Microenvironmental Control of Reindeer Antler Regeneration

**DOI:** 10.64898/2026.11.09.765262

**Authors:** Hanna Saari, Loan Tu Pham, Saeka Shimochi, Valentine Jarry, Mika Suzuki, Ilkka Syvänperä, Ilari Angervo, Tsuyoshi Kimura, Petriina Paturi, Kari Saikkonen, Pere Puigbò, Miho Nakamura

## Abstract

Deer antlers are the only mammalian appendages capable of complete periodic regeneration and therefore provide a natural model for investigating mechanisms of skeletal regeneration. However, current knowledge of the structural and molecular characteristics of the regenerative antler microenvironment remains incomplete. Here, we compared antler tip and antler base tissues from *Rangifer tarandus* (reindeer) with mature metacarpal cortical bone using structural, compositional, histological, and proteomic analyses. Antler tissues exhibited smaller and less crystalline apatite structures, elevated carbonate substitution, and higher organic matrix content compared with metacarpal bone, indicating an immature mineral phase. Antlers also exhibited lower bone mineral density, greater porosity, reduced bone volume fraction, and greater trabecular separation, particularly in central regions, consistent with a dynamic remodeling microenvironment. Histological examination revealed abundant active osteons and extensive osteoid seams in antler tissues, whereas metacarpal bone predominantly comprised mature lamellar bone with limited remodeling activity. Proteomic profiling identified distinct extracellular matrix signatures between antler and metacarpal tissues. Metacarpal bone was enriched in proteins associated with skeletal homeostasis, mechanotransduction, and bone remodeling, whereas antler tissues were enriched in proteins involved in extracellular matrix reorganization, metabolic adaptation, and regeneration. Collectively, these findings indicate that reindeer antlers possess a specialized regenerative skeletal microenvironment characterized by immature mineral organization, active remodeling, and distinct molecular signatures associated with rapid tissue growth. These results provide new insights into the biological basis of mammalian appendage regeneration and may facilitate the development of regenerative strategies for bone repair and skeletal tissue engineering.

## Introduction

Bone is a highly dynamic tissue that undergoes continuous remodeling to maintain structural integrity and metabolic balance ^1–3^. Although adult mammalian bone possesses substantial reparative capacity, complete organ-level regeneration is exceptionally rare. Deer antlers are a remarkable exception, representing the only mammalian appendage capable of complete annual regeneration ^4–7^. Antlers are rapidly growing temporary skeletal organs that are cast and regenerated each year through a process involving extensive angiogenesis, chondrogenesis, endochondral ossification, and mineralization. In reindeer (*Rangifer tarandus*), both males and females regenerate antlers, making this species a particularly valuable model for investigating regenerative skeletal biology ^8^.

During the growth phase, antler elongation rates can reach approximately 2–2.75 cm per day, making antlers among the fastest growing mineralized tissues in mammals ^7,9^. Despite this rapid growth, antlers maintain highly organized tissue architecture and controlled mineralization without malignant transformation. This unique balance between rapid proliferation and physiological regulation has generated increasing interest in antlers as a model for regenerative medicine, bone remodeling, and tissue engineering ^4–7,10^. On a systemic level, antler regeneration is closely linked to whole-body mineral metabolism. During antler growth, deer undergo a transient state of cyclic physiological osteoporosis, characterized by reversible skeletal demineralization followed by full restoration once antler mineralization is complete ^9^. This natural cycle models coupled bone loss and recovery, offering insights into mechanisms that could be leveraged to prevent or treat human osteoporosis.

At the tissue level, antlers possess a highly specialized skeletal microenvironment distinct from mature cortical bone. Antler tissue undergoes continuous matrix deposition, mineralization, vascular invasion, and osteonal remodeling throughout growth ^4–7,10^. These processes generate pronounced spatial heterogeneity in mineral density, trabecular organization, porosity, and extracellular matrix composition between the antler bone tissue and mature cortical bone ^7,11^. Omics-based studies further indicate that antler ossification involves pathways shared by vascular and neural regeneration, while transiently activating select cancer-associated signaling programs—highlighting the tightly regulated coordination required to achieve organized microarchitecture during rapid growth ^12–17^. Understanding these structural and compositional dynamics not only illuminates fundamental bone biology but also informs the development of regenerative microenvironments, mineralization strategies, and potential therapeutic targets for bone diseases.

Previous studies have shown that antler regeneration is mediated by specialized stem and progenitor cell populations located within the pedicle periosteum and reserve mesenchyme, including Paired Related Homeobox 1⁺ mesenchymal progenitors and antler blastema progenitor cells, which exhibit strong self-renewal and multilineage differentiation potential ^4,18,19^. Despite their extremely high rates of proliferation, antlers appear remarkably resistant to tumorigenesis. Mechanistic studies suggest that antlers mitigate oncogenic risk through enhanced apoptosis in the reserve mesenchyme, seasonal elimination of hyperproliferative tissue via casting, and positive selection of p53-associated regulatory genes ^20^. This unique intersection of rapid growth and cancer resistance positions antlers as an exceptional model for understanding how proliferative tissues maintain genomic stability and tissue-level homeostasis.

Despite growing interest in antler biology, the structural and molecular organization of the antler bone microenvironment remains incompletely understood. In particular, integrated analyses linking mineral organization, extracellular matrix composition, trabecular architecture, and proteomic signatures across different antler regions remain limited. Therefore, the present study aimed to characterize the regional microenvironmental differences between reindeer metacarpal and antler tissues, including antler tip and antler base. We quantified microarchitecture and mineralization, bioapatite crystal structure, and molecular composition. Multi-region quantitative analysis and standardized statistical testing were used to resolve site-specific gradients within antler tissue and establish comparative structural baselines relevant for regenerative biology and biomaterials research.

## Results

### Structural composition of bone and antler samples

The X-ray diffraction (XRD) patterns from all bone specimens closely matched the stoichiometric hydroxyapatite reference standard (HAp; ICDD 9-432), confirming hydroxyapatite as the predominant mineral phase in both antler and metacarpus tissues (**Fig. 1a**). Diffraction parameters derived from the (002) and (004) reflections were used to evaluate crystallographic properties, including crystallite size, d-spacing, and c-axis orientation (**Table 1**). For the (002) reflection, metacarpal samples exhibited narrower diffraction peaks (lower FWHM) than antler tissues, indicating larger crystallite size. Although metacarpus samples showed slightly higher mean 2θ values, the differences among the groups were small. Similar results were observed for the (004) reflection, with metacarpus exhibiting narrower diffraction peaks than antler tissues. In contrast, d-spacing values showed little variation among the three groups, particularly for the (004) reflection. Overall, Scherrer analysis indicated that antler tissues contained smaller crystallites than metacarpus, consistent with a less mature apatite crystal structure.

**Fig. 1.**
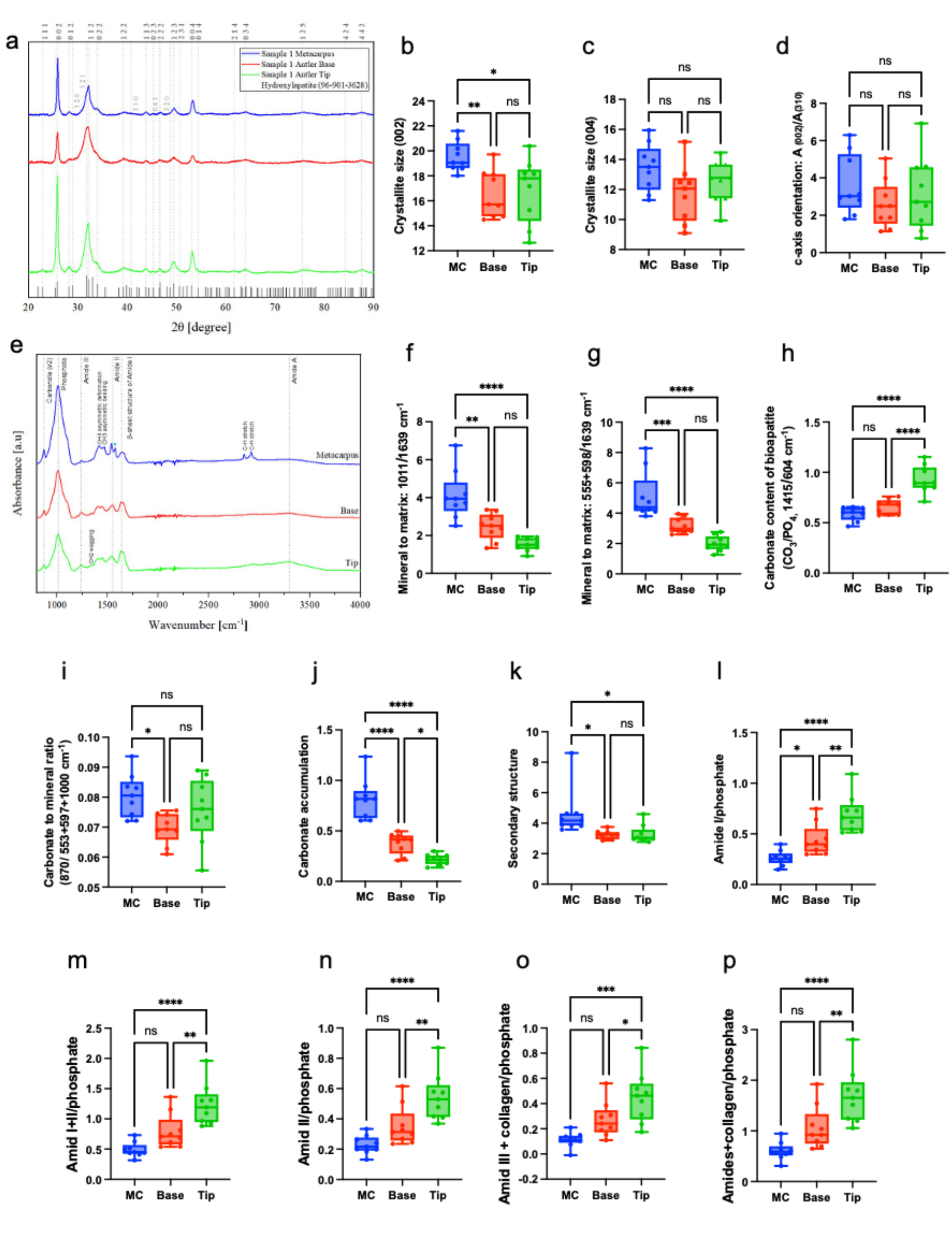
XRD and ATR-FTIR characterization of reindeer metacarpal and antler bone tissues. **(a)** XRD patterns of metacarpus (MC), antler base, and antler tip compared with the stoichiometric hydroxyapatite reference standard, confirming single-phase hexagonal hydroxyapatite as the major mineral phase in all tissues. **(b-c)** Crystallite size derived from the (002) and (004) reflections using the Scherrer equation. Metacarpus exhibits larger crystallite size than antler tissues. **(d)** Preferred c-axis orientation expressed as the A₍₀₀₂₎/A₍₃₁₀₎ area ratio. No statistically significant differences were observed among groups. **(e)** Representative ATR-FTIR spectra showing characteristic phosphate, carbonate, and collagen-associated absorption bands. **(f-g)** Mineral-to-matrix ratios showing significantly higher mineralization in metacarpus compared with antler tissues. **(h-j)** Carbonate-related parameters indicating differences in carbonate substitution and accumulation among tissues. **(k-p)** Protein-related FTIR parameters associated with collagen and secondary protein structure. Antler tip showed elevated amide-related ratios, whereas metacarpus exhibited stronger overall protein-associated signals. Data are presented as the mean ± SD (n=9). Statistical significance was determined using one-way ANOVA with Tukey’s HSD post hoc test. * *p* < 0.05, \*\**p* < 0.002, \*\*\**p* < 0.001, \*\*\*\**p* < 0.0001.

**Table 1.** XRD-derived structural parameters for metacarpal, antler tip, and antler base bone tissues. Reported values include 2θ peak positions, full width at half maximum (FWHM), and calculated interplanar spacings (d) for the (002) and (004) reflections. FWHM and d-spacing values were obtained from peaks refined using the Nonlinear Fit (NLFit) function. Values are presented as mean ± SD (n=9).

| Miller indices (hkl) | Sample type | 2θ [°] | FWHM | d [Å] |
|---|---|---|---|---|
| 002 | Metacarpus | 25.88 ± 0.06 | 0.44 ± 0.03 | 3.44 ± 0.008 |
| 002 | Antler tip | 25.79 ± 0.11 | 0.52 ± 0.09 | 3.45 ± 0.01 |
| 002 | Antler base | 25.79 ± 0.12 | 0.52 ± 0.06 | 3.45 ± 0.02 |
| 004 | Metacarpus | 53.25 ± 0.07 | 0.70 ± 0.08 | 1.72 ± 0.002 |
| 004 | Antler tip | 53.19 ± 0.11 | 0.75 ± 0.09 | 1.72 ± 0.003 |
| 004 | Antler base | 53.20 ± 0.11 | 0.80 ± 0.13 | 1.72 ± 0.003 |

Crystallite size calculated using the Scherrer equation revealed significantly larger crystallites in metacarpus compared with antler tissues for the (002) reflection (**Fig. 1b**). Metacarpal samples exhibited the largest mean crystallite size, whereas antler tip and base displayed smaller and similar values. No significant differences were observed among groups for crystallite size derived from the (004) reflection (**Fig. 1c**). The c-axis orientation index, calculated as the area ratio of the (002) and (310) (A₍₀₀₂₎/A₍₃₁₀₎) was highest in metacarpus and lowest in antler base; however, these differences did not reach statistical significance (**Fig. 1d**). Collectively, these findings indicate that metacarpus possesses a more crystalline mineral structure than antler tissue.

The attenuated total reflectance Fourier transform infrared spectroscopy (ATR-FTIR) spectra from all bone samples exhibited characteristic features of mineralized collagenous tissue, including phosphate, carbonate, and collagen-associated absorption bands (**Fig. 1e**) ^21–23 24^. Major phosphate vibrations were observed at ∼1011 cm⁻¹ with associated shoulders near 960 cm⁻¹ and 1103 cm⁻¹, confirming hydroxyapatite as the dominant mineral phase. Collagen-associated amide I, II, and III bands were detected at ∼1639, 1546, and 1239 cm⁻¹, respectively. Carbonate substitution within the apatite lattice was indicated by the ν_2_ carbonate band at ∼871 cm⁻¹.

Mineral-to-matrix ratios calculated using both phosphate/amide I ^25^ and phosphate splitting-factor ^23^ approaches demonstrated significantly greater mineralization in metacarpus compared with antler tissues (**Fig. 1f-g**). No significant differences were observed between antler tip and base.

Carbonate-related parameters demonstrated distinct tissue-specific trends. The CO₃/PO₄ ratio ^21^ was highest in antler tip tissue, indicating increased carbonate substitution in rapidly growing antler (**Fig. 1h**). In contrast, both the carbonate-to-mineral ratio and carbonate accumulation index were significantly higher in metacarpus compared with antler tissues (**Fig. 1i–j**) ^23^, suggesting greater structural carbonate incorporation in mature cortical bone.

Protein secondary structure signals were significantly greater in metacarpus than in antler tissues (**Fig. 1k**). In contrast, all calculated amide/phosphate-related parameters were elevated in antler tissues, particularly in antler tip specimens (**Fig. 1l–p**), indicating a greater organic matrix contribution in actively growing antler tissue.

Micro-computed tomography (micro-CT) analysis revealed clear regional differences in bone mineral density (BMD) among metacarpus, antler base, and antler tip samples (**Fig. 2a–b**). Metacarpus exhibited the highest overall BMD (1.0007 g/cm³ ± 0.064), with low inter-individual variation. In both antler tip and antler base, BMD was lowest in the central region and increased toward the intermediate and peripheral regions. Total and open porosity differed significantly among metacarpus and the regional compartments of antler tip and antler base tissues (**Fig. 2c–d**). The central regions of both antler tip and antler base exhibited markedly greater porosity than the corresponding intermediate and peripheral regions, whereas metacarpus displayed minimal porosity overall. Closed porosity showed no significant differences among the analyzed regions (**Fig. 2e**).

**Fig. 2.**
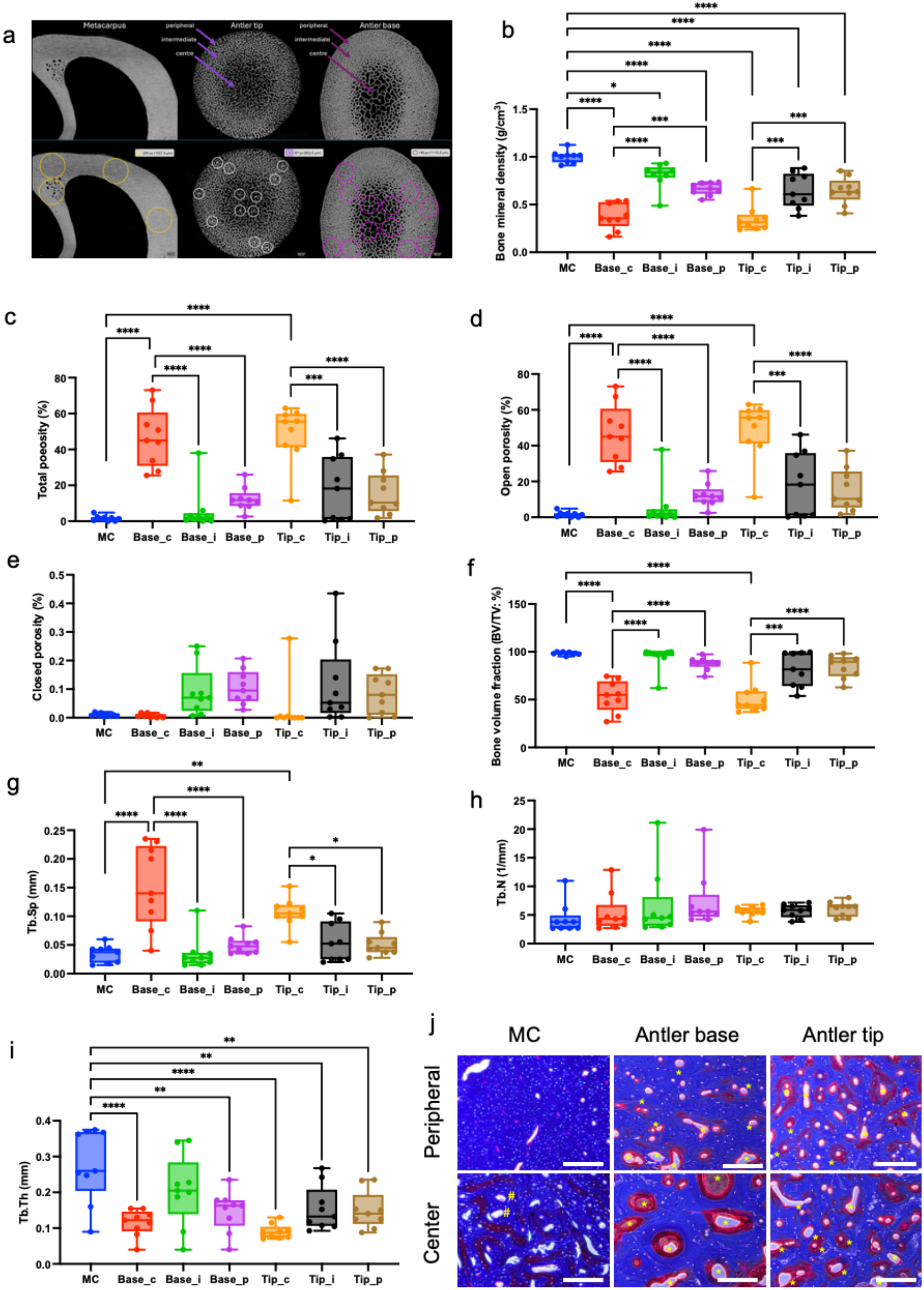
Micro-CT and histological analysis of reindeer metacarpal and antler bone tissues. **(a)** Representative micro-CT cross-sectional images and ROI locations in metacarpus, antler base, and antler tip. Circular ROIs were positioned within peripheral (_p), intermediate (_i), and central (_c) regions of antler tissues, and within cortical regions of the metacarpus. **(b)** BMD analysis showed highest and most uniform mineral density in metacarpus, while antler tissues exhibited lower BMD with regional variation increasing toward the periphery. **(c-e)** Total, open, and closed porosity measurements. Antler central regions showed the highest open and total porosity, whereas metacarpus exhibited the lowest porosity overall. Closed porosity did not differ significantly among groups. **(f)** Bone volume fraction (BV/TV), with highest values in intermediate antler regions and lowest values in central regions. **(g)** Trabecular separation (Tb.Sp), showing highest values in central antler regions and lowest values in peripheral regions. **(h)** Trabecular number (Tb.N), showing no significant regional differences. **(i)** Trabecular thickness (Tb.Th), which was greater in intermediate antler regions. **(j)** Masson’s trichrome staining of reindeer metacarpus, antler base, and antler tip showing differences in bone matrix maturity and osteonal activity. Metacarpus exhibited predominantly mature cortical bone matrix with quiescent osteons (#), whereas antler tissues contained both mature and immature bone matrix with abundant active osteons (*) and thick osteoid seams throughout the tissue. Scale bars: 100 µm. Data are presented as the mean ± SD (n=9). Statistical significance was determined using one-way ANOVA with Tukey’s HSD post hoc test. * *p* < 0.05, \*\**p* < 0.002, \*\*\**p* < 0.001, \*\*\*\**p* < 0.0001.

Bone volume fraction (BV/TV) differed significantly among metacarpus and central regions of antler tissues (**Fig. 2f**). In antler tissues, the central regions of both antler tip and antler base demonstrated the lowest BV/TV values, indicating reduced mineralized bone volume within the inner antler core. Trabecular separation (Tb.Sp) differed significantly among metacarpus, antler tip, and antler base regions (**Fig. 2g**). Metacarpus exhibited the lowest apparent Tb.Sp values, reflecting its dense cortical organization and limited internal porosity. In contrast, the center of the antler base demonstrated the highest Tb.Sp values, indicating the most open trabecular architecture among all analyzed regions. Trabecular thickness (Tb.Th) differed significantly among metacarpus, antler tip, and antler base regions (**Fig. 2i**). Metacarpus exhibited greater structural thickness values, consistent with its dense cortical organization. Trabecular number (Tb.N) showed no statistically significant differences among metacarpus, antler tip, and antler base regions (**Fig. 2h**). These findings indicate that antler tissues exhibit a progressively more compact and mineralized architecture from the central to peripheral regions, whereas metacarpal bone retains the densest and least porous microstructure overall.

### Histological analysis of bone and antler samples

Masson’s trichrome staining revealed clear differences in tissue organization between metacarpal and antler samples (**Fig. 2j**). In metacarpus, the peripheral region of the bone cross-sections showed a dense, mature bone matrix, evidenced by strong blue staining. In contrast, the central region displayed a predominantly immature bone matrix, stained red–purple. Most osteons within the metacarpal mid-region were identified as quiescent osteons, whereas fewer were observed near the outer cortical area.

Both antler base and antler tip contained mixtures of mature and immature bone matrix throughout their cross-sections. Active osteons, characterized by red-stained osteoid, were present in both peripheral and central regions. Compared with metacarpus, antler samples exhibited a higher number of active osteons, with notably thicker osteoid seams particularly prominent in the central regions of the cross-sections.

### Proteomics analysis of bone and antler samples

Proteomic analysis identified distinct molecular signatures among metacarpus, antler base, and antler tip samples (**Fig. 3, Tables S1-S7**). Based on statistically significant differential protein expression patterns, proteins were categorized into eight groups (**Fig. 3a**): Group i, proteins distinguishing metacarpus from antler tip and base (n = 16); Group ii, proteins distinguishing antler tip from metacarpus and base (n = 4); Group iii, proteins distinguishing antler base from metacarpus and tip (n = 10); Group iv, proteins shared between metacarpus and antler base (n = 13); Group v, proteins shared between antler tip and base (n = 2); Group vi, proteins shared between metacarpus and antler tip (n = 7); Group vii, proteins differentially expressed among all three tissues (n = 1); and Group viii, proteins similarly expressed across metacarpus, antler tip, and antler base (n = 162).

**Fig. 3.**
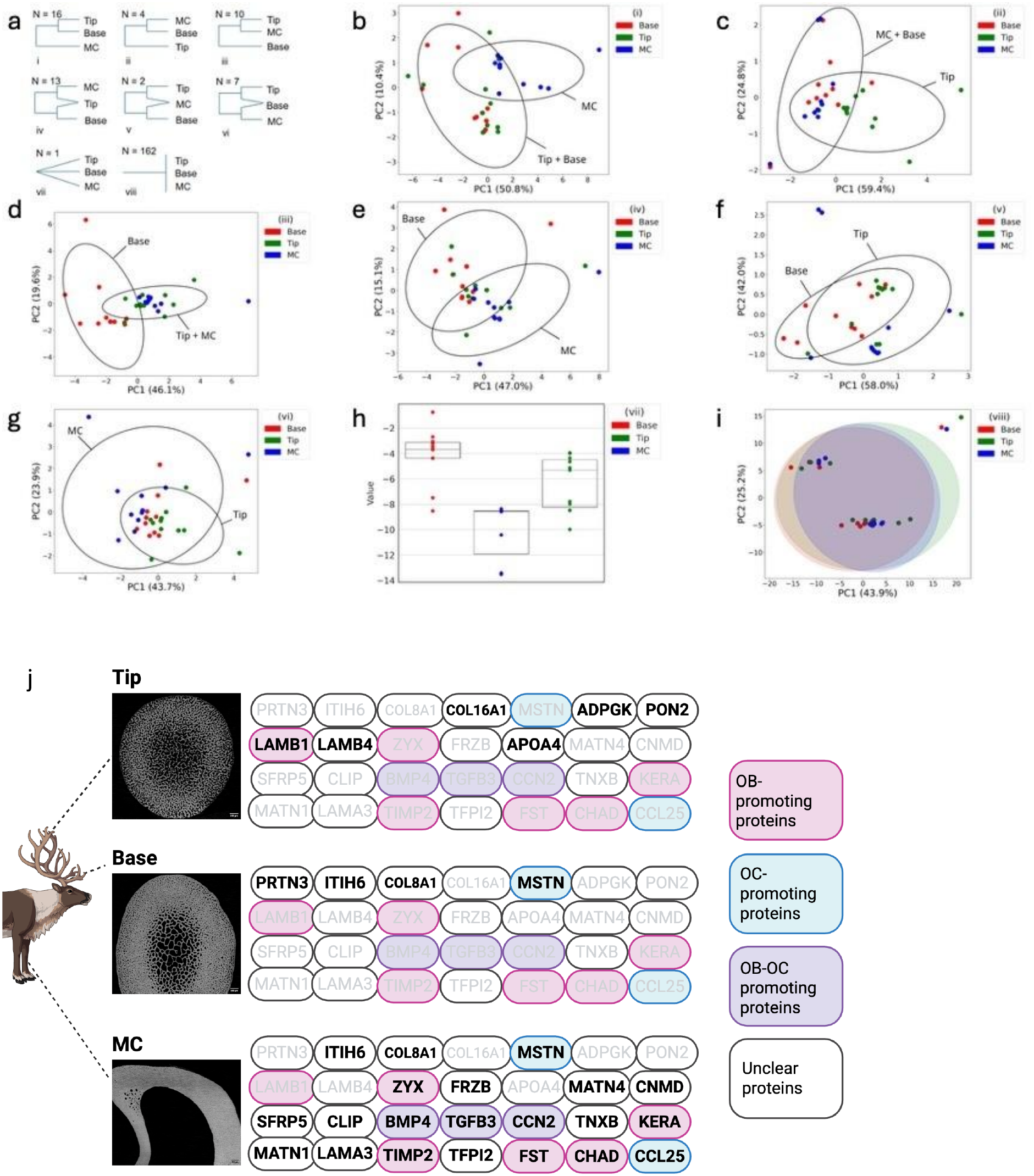
Proteomics analysis of reindeer bone samples. T-test results and visualization of t-test based groups. Colors represent different sample types Tip, Base and Metacarpus (MC). **(a)** Proteins groups based on t-test results, **(b)** PCA plot of group i consisting of proteins that separate metacarpus from tip and base, **(c)** PCA plot of group ii consisting of proteins that separate tip from metacarpus and base, **(d)** PCA plot of group iii consisting of proteins that separate base from metacarpus and tip, **(e)** PCA plot of group iv consisting of proteins that separate metacarpus from base, **(f)** PCA plot of group v consisting of proteins that separate base from tip, **(g)** PCA plot of group vi consisting of proteins that separate metacarpus from tip, **(h)** Box and scatter plot of group vii consisting of proteins that separate tip, base, and metacarpus from each other, **(i)** PCA plot of group viii consisting of proteins that appear similarly in all base, tip, and metacarpus. **(j)** Schematic summary of representative ECM-associated proteins enriched in each tissue type. Metacarpus showed enrichment of 18 ECM proteins, including ZYX, FRZB, MATN4, CNMD, SFRP5, CILP, BMP4, TGFB3, CCN2, TNXB, KERA, MATN1, LAMA3, TIMP2, TFPI2, FST, CHAD, and CCL25. Antler tip tissue preferentially expresses 5 proteins, including ADPGK, PON2, LAMB1, LAMB4, and APOA4. Protein label colors indicate their reported biological functions in bone remodeling: red, osteoblast (OB)-promoting proteins; blue, osteoclast (OC)-promoting proteins; violet, proteins associated with both osteoblast and osteoclast (OB-OC) promotion; and white, proteins with unclear or insufficiently characterized dominant functions. Created in BioRender. Nakamura, M. (2026) https://BioRender.com/dmjivo1

Principal component analysis (PCA) shows clear clustering patterns among metacarpus, antler base, and antler tip specimens, indicating tissue-specific protein expression profiles (**Fig. 3b–g**). In Groups i–iii, the covariance ellipses exhibited distinct orientations and distributions, suggesting differences in proteomic organization and tissue-specific molecular composition among the three skeletal tissues. The box-and-scatter plot of Group vii (**Fig. 3h**) demonstrated a distinct distribution pattern among metacarpus, antler base, and antler tip, with antler base showing overall higher protein abundance than the other groups. In contrast, PCA analysis of Group viii revealed substantial overlap among covariance ellipses (**Fig. 3i**), indicating similar proteomic variability and shared baseline protein expression across all tissues.

Several extracellular matrix (ECM) proteins related to mature bone tissue, including zyxin (ZYX), Secreted frizzled-related protein 3 (FRZB), Matrilin-4 (MATN4), Chondromodulin (CNMD), Secreted frizzled-related protein 5 (SFRP5), CAP-Gly Cartilage intermediate layer protein 1 (CILP) Bone morphogenetic protein 4 (BMP4), Transforming growth factor β3 (TGFB3), Cellular communication network factor 2 (CCN2), Tenascin-X (TNXB), Keratocan (KERA), Matrilin-1 (MATN1), Laminin subunit alpha-3 (LAMA3), Tissue inhibitor of metalloproteinases 2 (TIMP2), Tissue factor pathway inhibitor 2 (TFPI2), Follistatin (FST), Chondroadherin (CHAD), C-C motif chemokine ligand 25 (CCL25) were markedly enriched in metacarpus, whereas antler bone tissues exhibited comparatively lower expression of these mature bone-associated proteins (**Table 2 and Fig. 3j**). In contrast, proteins associated with tissue remodeling and regeneration, including ADP-dependent glucokinase (ADPGK), Paraoxonase 2 (PON2), Laminin subunit beta-1 (LAMB1), Laminin subunit beta-4 (LAMB4), and Apolipoprotein A-IV (APOA4), were preferentially enriched in antler tip specimens. Conversely, myostatin (MSTN), inter-alpha-trypsin inhibitor heavy chain 6 (ITIH6), and collagen alpha-1(VIII) chain (COL8A1) exhibited lower abundance in antler tip tissue, whereas proteinase 3 (PRTN3) was preferentially enriched in antler base tissue. Overall, proteins associated with tissue remodeling, development, and regeneration were more abundant in antler tissues, supporting the concept that antlers represent a continuously regenerating skeletal tissue with biological properties distinct from those of conventional cortical bone such as the metacarpus. Distinct protein expression profiles were observed between metacarpal bone and antler tissues.

**Table 2.** List of extracellular matrix (ECM) proteins that present significant differences (p-value < 0.05) among reindeer’s antler (tip, base) and metacarpal tissues. A full version of this table is available in supplementary Table S8.

| Protein | MC | AT-Base | AT-Tip | Phylobone Code | Type | OB | OC |
|---|---|---|---|---|---|---|---|
| PRTN3 | - | + | n.s | PB0014 | Putative regulatory | n.e. | n.e. |
| ITIH6 | + | + | - | PB0093 | Other/Unknown | n.e. | n.e. |
| COL8A1 | + | + | - | PB0040 | Structural | n.e. | n.e. |
| COL16A1 | - | - | + | PB0091 | Structural | n.e. | n.e. |
| MSTN | + | + | - | PB0178 | Other/Unknown | n.e. | ↑ |
| ADPGK | - | n.s | + | PB0020 | Other/Unknown | n.e. | n.e. |
| PON2 | - | n.s | + | PB0207 | Other/Unknown | n.e. | n.e. |
| LAMB1 | - | n.s | + | PB0128 | Structural | ↑ | n.e. |
| LAMB4 | - | n.s | + | PB0105 | Structural | n.e. | n.e. |
| ZYX | + | n.s | - | PB0244 | Other/Unknown | ↑ | n.e. |
| FRZB | + | n.s | - | PB0220 | Regulatory | n.e. | n.e. |
| APOA4 | - | n.s | + | PB0049 | Other/Unknown | n.e. | n.e. |
| MATN4 | + | - |  | PB0194 | Other/Unknown | n.e. | n.e. |
| CNMD | + | - |  | PB0039 | Regulatory | n.e. | n.e. |
| SFRP5 | + | - |  | PB0119 | Other/Unknown | n.e. | n.e. |
| CILP | + | - |  | PB0099 | Structural | n.e. | n.e. |
| BMP4 | + | - |  | PB0251 | Regulatory | ↑ | ↑ |
| TGFB3 | + | - |  | PB0201 | Regulatory | ↑ | ↑ |
| CCN2 | + | - |  | PB0196 | Regulatory | ↑ | ↑ |
| TNXB | + | - |  | PB0063 | Structural | n.e. | ↓ |
| KERA | + | - |  | PB0197 | Other/Unknown | ↑ | n.e. |
| MATN1 | + | - |  | PB0031 | Structural & Regulatory | n.e. | n.e. |
| LAMA3 | + | - |  | PB0135 | Structural | n.e. | n.e. |
| TIMP2 | + | - |  | PB0188 | Other/Unknown | ↑ | ↓ |
| TFPI2 | + | - |  | PB0042 | Structural | n.e. | ↓ |
| FST | + | - |  | PB0221 | Regulatory | ↑ | n.e. |
| CHAD | + | - |  | PB0007 | Regulatory | ↑ | ↓ |
| CCL25 | + | - |  | PB0106 | Other/Unknown | n.e. | ↑ |

MC: Metacarpal, AT: Antler, OB: Osteoblast, OC: Osteoclast, +: *higher expression,* -: *lower expression, n.s.: not significant,* ↑: promotion,↓: inhibition, n.e.: no evidence.

### Multilineage Differentiation of Reindeer Mesenchymal Stem Cells

Reindeer mesenchymal stem cells (MSCs) demonstrated the capacity to differentiate into osteogenic, adipogenic, and chondrogenic lineages under lineage-specific induction conditions (**Supplementary Fig. S2**). Following osteogenic stimulation, cultures showed prominent calcium-rich mineral deposits that stained intensely red with Alizarin Red S, confirming successful osteoblastic differentiation. These mineralized nodules were distributed throughout the culture, indicating extracellular matrix mineralization.

Adipogenic induction resulted in the appearance of intracellular lipid droplets, which were positively stained using Oil Red O. These lipid inclusions, visible as distinct red-brown vesicles within the cytoplasm, confirmed adipocyte formation and lipid accumulation characteristic of terminal adipogenic differentiation. Under chondrogenic conditions, cells formed dense, rounded aggregates typical of cartilaginous micromass cultures. Histological analysis revealed a strongly basophilic extracellular matrix consistent with proteoglycan-rich cartilage tissue, confirming successful differentiation into chondrocytes. Together, these findings verify the multilineage differentiation potential of reindeer MSCs, demonstrating their ability to generate bone, fat, and cartilage cell types under lineage-specific induction.

## Discussion

### Skeletal microenvironment in the antler as key differential factor

The present study demonstrates that reindeer antler tissue possesses a highly specialized skeletal microenvironment that differs substantially from mature cortical bone such as the metacarpus. By integrating crystallographic, spectroscopic, microarchitectural, histological, and proteomic analyses, our findings suggest that antler tissue represents a dynamic regenerative microenvironment optimized for rapid growth, active remodeling, and continuous matrix turnover ^6,18,26,27^. In contrast, metacarpus exhibited structural and molecular features characteristic of mature load-bearing cortical bone with relatively stable mineral homeostasis ^28,29^. Antlers are unique among mammalian organs because they undergo complete periodic regeneration through rapid endochondral ossification and extensive extracellular matrix remodeling, making them an important model for studying skeletal regeneration and regenerative osteogenesis ^26,27^.

A prominent feature distinguishing antler and metacarpus tissues was their mineral organization and matrix composition. XRD analysis demonstrated that antler tissues contained smaller and less crystalline apatite crystallites than metacarpus, particularly along the (002) plane. Reduced crystallite size and broader diffraction peaks are generally associated with immature or newly deposited mineral phases, suggesting that antler tissue undergoes continuous mineral deposition and remodeling ^30,31^. The lower mineral maturity observed in antler tissues is consistent with the extraordinary growth rate of deer antlers, which can exceed several centimeters per day during peak growth periods ^26,32^. In contrast, metacarpus exhibited larger and more ordered apatite crystals, reflecting long-term mineral maturation under sustained mechanical loading conditions ^28,33^.

ATR-FTIR analysis further supported the existence of a distinct antler microenvironment characterized by elevated organic matrix content and altered carbonate incorporation. Antler tissues, particularly the antler tip, exhibited higher amide/phosphate-related ratios and increased carbonate substitution compared with metacarpus. These findings indicate that antler tissue contains a less mineralized but more protein-rich extracellular matrix, consistent with active osteoid formation and rapid collagen synthesis ^25,34,35^. In contrast, metacarpus showed greater mineral-to-matrix ratios and higher structural carbonate accumulation, which are hallmarks of mature cortical bone ^30,36^. The coexistence of lower crystallinity, higher organic matrix content, and elevated carbonate substitution strongly suggests that the antler microenvironment remains in a relatively immature and metabolically active state throughout growth ^6,25^. The apparent discrepancy between these carbonate metrics reflects the fact that they quantify different aspects of carbonate incorporation and mineral maturity.

Micro-CT and histological analyses revealed profound differences in tissue architecture that likely contribute to the regenerative capacity of antlers. Metacarpal bone displayed a dense cortical structure characterized by high mineral density, low porosity, and limited internal spacing between mineralized structures. In contrast, antler tissues exhibited lower BMD, markedly increased porosity, and greater trabecular separation, particularly within the central regions. These structural characteristics likely facilitate rapid vascular invasion, nutrient diffusion, and cellular migration during antler elongation and mineralization ^26,27,32^. Previous studies have demonstrated that growing antlers contain extensive vascular networks and highly active remodeling zones that support rapid tissue expansion and endochondral ossification ^27,32^. The central antler regions demonstrated the most porous architecture and lowest bone volume fraction, whereas peripheral regions exhibited progressive structural compaction and mineral accumulation. Such spatial heterogeneity suggests the presence of region-specific microenvironments within antlers that correspond to different stages of tissue maturation, mineralization, and remodeling^28,32,37^.

Histological findings further emphasized the highly active remodeling state of antler tissue. Masson’s trichrome staining demonstrated abundant active osteons and thick osteoid seams throughout both antler base and antler tip, whereas metacarpus predominantly contained quiescent osteons embedded within mature lamellar bone. The coexistence of immature and mature matrix components throughout antler tissue indicates continuous bone formation and remodeling rather than the relatively balanced turnover observed in mature cortical bone ^26,38^. Importantly, the active osteonal remodeling observed in antlers resembles certain aspects of fracture healing and developmental osteogenesis, suggesting that the antler microenvironment may recapitulate regenerative pathways that are otherwise restricted in adult mammalian skeletal tissues ^27,39,40^. Previous studies have similarly described antlers as a unique example of physiologically regulated mammalian organ regeneration involving rapid endochondral ossification and extensive osteogenic activity ^26,27^ .

### Proteomic signatures of the regenerative antler microenvironment

The observed microenvironmental differences between metacarpal and antler tissues likely reflect their fundamentally different biological functions. Metacarpus functions primarily as a mechanically stable load-bearing organ that prioritizes structural rigidity and mineral maturity ^33,41^. In contrast, antlers are transient skeletal appendages that undergo rapid seasonal regeneration and therefore require a highly plastic microenvironment capable of supporting intense cellular proliferation, vascularization, matrix remodeling, and mineral deposition ^7,26,27^. Previous studies have shown that antler growth is accompanied by extensive angiogenesis, rapid endochondral ossification, and high metabolic activity, making antlers one of the fastest growing mammalian tissues ^7,26,27^. The elevated porosity, immature mineral phase, active osteoid formation, and regenerative proteomic profile identified in antlers collectively support this interpretation and further emphasize the unique regenerative biology of antler tissue ^7,10^.

Proteomics analysis provided additional insight into the molecular mechanisms underlying the specialized antler microenvironment. PCA showed clear separation between metacarpal and antler tissues, supporting the presence of distinct structural and regulatory protein signatures (**Fig. 3b-i**). Metacarpus was enriched in several proteins involved in skeletal matrix organization, bone remodeling, mechanotransduction, and maintenance of tissue homeostasis, including CCN2, ZYX, FRZB, MATN4, CNMD, SFRP5, BMP4, TGFB3, TNXB, TIMP2, FST, CHAD, and CCL25 (**Fig. 3j**). Many non-collagenous extracellular matrix proteins are increasingly recognized as critical regulators of bone remodeling, matrix organization, cell–matrix interactions, and mechanosensitive signaling pathways that contribute to skeletal integrity and adaptation to mechanical loading ^42,43^. In particular, CCN2 has been shown to regulate osteoblast differentiation and osteoclastogenesis while promoting extracellular matrix production and skeletal remodeling ^44,45^. Similarly, BMP4 and TGFB3 are well-established regulators of osteogenesis, skeletal development, and bone remodeling, whereas FST modulates activin and myostatin signaling pathways involved in bone formation and resorption. ZYX functions as a mechanosensitive focal adhesion protein involved in cytoskeletal organization and cellular responses to mechanical loading ^46,47^. The enrichment of these proteins in metacarpus therefore likely reflects the highly organized remodeling processes and continuous adaptation to weight-bearing mechanical stress characteristic of mature cortical bone.

In contrast, antler tissues exhibited enrichment of proteins associated with extracellular matrix remodeling, cellular metabolism, tissue regeneration, and developmental plasticity, including APOA4, LAMB1, LAMB4, PON2, and ADPGK (**Fig. 3j**). Several of these proteins have been linked to cellular adaptation, migration, metabolic reprogramming, and tissue remodeling in rapidly growing or regenerating tissues. The enrichment of APOA4 in antler tip tissue is particularly noteworthy. Although APOA4 has been reported as a biomarker in several cancers and proliferative conditions ^48–50^, current evidence does not strongly support a direct role in tumor progression. Instead, its elevated abundance in antler tissue may reflect increased metabolic activity, lipid transport, and oxidative stress regulation associated with rapid tissue growth. Similarly, PON2 has been implicated in cellular protection against oxidative stress and metabolic adaptation ^51^, whereas ADPGK is involved in alternative glucose metabolism that may support the energetic demands of rapidly proliferating tissues ^52^. Although these associations do not establish direct regenerative functions for these proteins in antlers, their enrichment is consistent with the elevated metabolic and biosynthetic activity required to sustain rapid antler growth. Notably, MSTN expression was reduced in antler tip tissue but relatively enriched in antler base tissue and metacarpus (**Fig. 3j**). Given its established role as a negative regulator of musculoskeletal growth ^53,54^, this expression pattern may reflect regional differences in growth regulation and tissue maturation. Although the precise role of MSTN in antler biology remains unclear, its distribution suggests potential involvement in balancing skeletal growth and remodeling during the antler regeneration cycle.

### A Working Hypothesis on the regenerative microenvironment and skeletal progenitor cells

The multilineage differentiation capacity of reindeer metacarpus marrow-derived MSCs demonstrated in this study supports the presence of multipotent skeletal progenitor cells within the reindeer skeletal system. Although these cells were isolated from metacarpus rather than antler tissue, their differentiation potential is consistent with previous reports describing progenitor populations involved in antler regeneration ^6,27,57^. Under lineage-specific induction conditions, MSCs successfully differentiated into osteogenic, adipogenic, and chondrogenic lineages, demonstrating the capacity of reindeer skeletal progenitors to generate multiple mesenchymal cell types and contribute to skeletal maintenance and repair ^58,59^. Previous studies have identified stem cell-like populations within the pedicle periosteum and reserve mesenchyme of deer antlers, where they contribute to rapid antler regeneration through extensive proliferation and multilineage differentiation ^6,27,57^. Together, our findings and previous studies support the concept that cervids possess specialized progenitor cell populations that facilitate both routine skeletal remodeling and the remarkable regenerative capacity of antler tissues.

Importantly, antlers represent a valuable model for investigating both regenerative and pathological bone remodeling processes relevant to human health (**Fig. 4**). The antler microenvironment combines rapid bone formation, extensive angiogenesis, active extracellular matrix remodeling, and tightly regulated cellular proliferation without malignant transformation ^7,26,27^. Understanding how antler tissues coordinate these processes may provide insights into mechanisms underlying fracture repair, osteoporosis, bone tissue engineering, and skeletal regenerative medicine ^7,10,26,27^. In particular, the coordinated interactions among extracellular matrix proteins, growth factors, and skeletal progenitor cells that support antler regeneration remain incompletely understood and warrant further investigation ^60,61^. Elucidating these mechanisms may contribute to the development of novel regenerative strategies for enhancing bone repair and skeletal tissue regeneration.

**Fig. 4.**
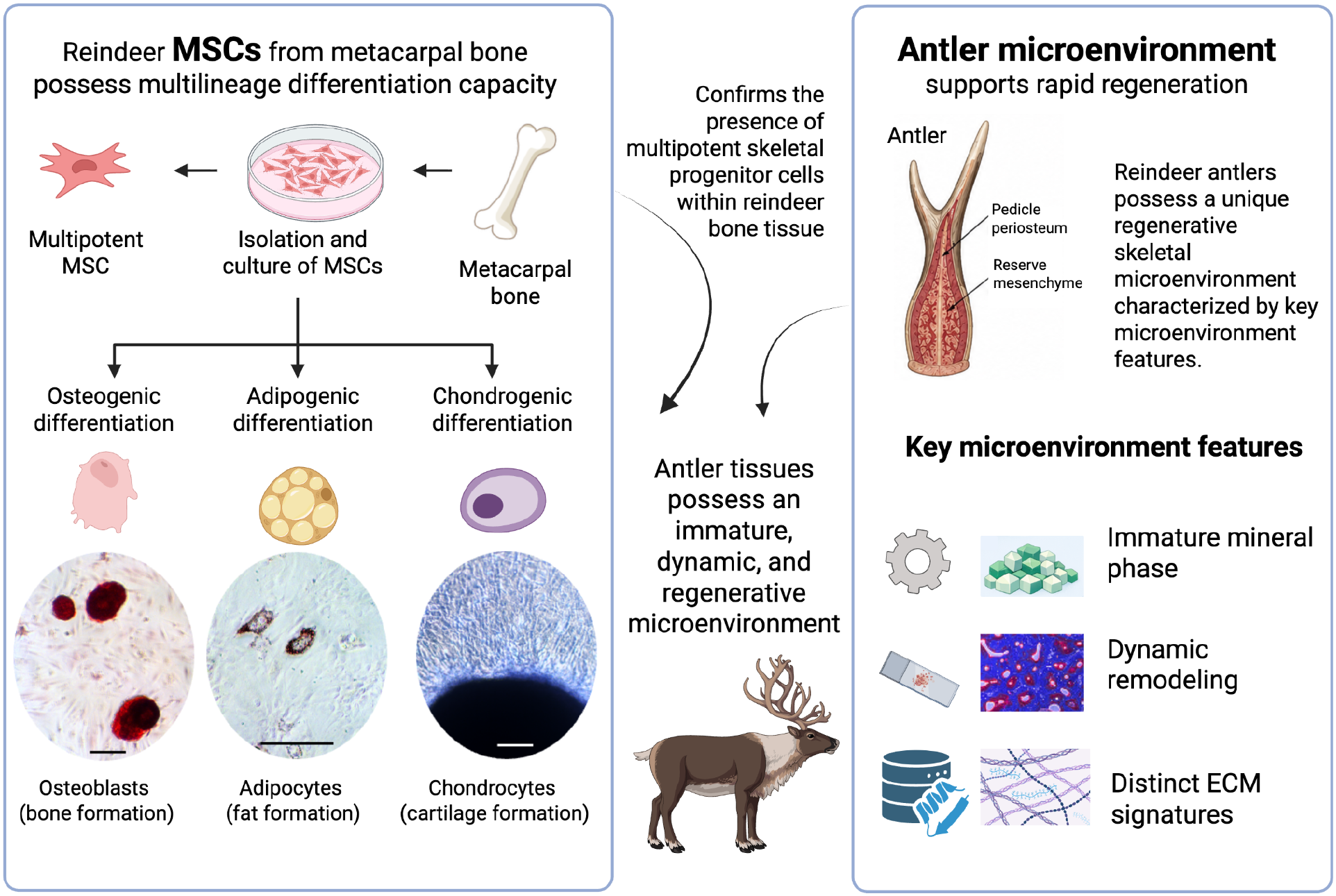
Proposed model of the regenerative skeletal microenvironment in reindeer antlers. Reindeer MSCs isolated from metacarpus exhibited multilineage differentiation capacity, as demonstrated by successful osteogenic, adipogenic, and chondrogenic differentiation (**Fig. S2**). The presence of multipotent skeletal progenitor cells supports the regenerative potential of reindeer skeletal tissues. Integrated structural, compositional, histological, and proteomic analyses revealed that antler tissues possess a distinct regenerative microenvironment compared with mature cortical bone. Antler tissues are characterized by an immature mineral phase with reduced crystallinity and increased organic matrix content, active tissue remodeling evidenced by abundant osteons and osteoid formation, and unique ECM protein signatures associated with regeneration and matrix turnover. Together, these findings support the concept that reindeer antlers contain a highly dynamic and regenerative skeletal microenvironment that facilitates rapid tissue growth, remodeling, and annual organ regeneration. Created in BioRender. Nakamura, M. (2026) https://BioRender.com/dmjivo1

Furthermore, the enrichment of laminin-related proteins, particularly LAMB1 and potentially LAMB4, suggests active remodeling of the extracellular matrix niche that may support progenitor cell maintenance, migration, and differentiation during antler regeneration ^62^. Laminins are major components of basement membrane-associated extracellular matrices ^63^ and regulate cell adhesion, migration, survival, and stem cell behavior during tissue development and regeneration ^62^. In addition, the relative enrichment of MSTN in antler base tissue, together with its reduced abundance in antler tip tissue, suggests regional differences in growth regulation and skeletal remodeling during the antler regeneration cycle. Given the established role of MSTN in musculoskeletal development and bone remodeling ^53,54^, this expression pattern may indicate involvement in the transition from active tissue growth to tissue maturation and resorption. However, the precise role of MSTN in antler biology remains unclear and requires further investigation. Together, these findings indicate that antler tissues maintain a regenerative microenvironment characterized by enhanced ECM turnover, metabolic activity, and regenerative plasticity, in contrast to the homeostatic and mechanically adapted protein profile observed in mature metacarpal tissue. These proteomic differences further support the concept that antlers represent a specialized regenerative skeletal tissue with biological properties distinct from those of conventional cortical bone.

### Limitations of the study

Antler tissues undergo dynamic seasonal and developmental changes, and the present analysis represents a single time point within the growth cycle. Previous studies have demonstrated substantial temporal variation in antler mineralization, vascularization, and remodeling activity throughout the regenerative cycle ^7,26,27^. Future longitudinal studies evaluating temporal changes in antler microenvironment composition may therefore provide additional insight into the regulation of regeneration and mineralization. Moreover, although proteomic analyses identified several candidate proteins associated with regeneration and mechanosensing, functional validation of these pathways remains necessary. Furthermore, it should be noted that these functions are largely inferred from human and mouse studies, and that proteins in reindeer may play alternative roles.

### Concluding remarks

The present study indicates that reindeer antler tissue possesses a unique regenerative skeletal microenvironment distinct from mature cortical bone. Antler tissues are characterized by immature mineral organization, increased organic matrix content, elevated porosity, active osteonal remodeling, and enrichment of regenerative extracellular matrix proteins. These integrated structural and molecular features likely facilitate the remarkable regenerative capacity of antlers and highlight their value as a model system for studying skeletal regeneration, extracellular matrix biology, and bone remodeling.

## Materials and Methods

### Sample preparation

Reindeer samples were obtained from the Kaldoaivi reindeer herding cooperative at Skalluvaara, Utsjoki, Finland (69°80′N, 27°17′E), and processed at the Kevo Research Station, Department of Biodiversity Sciences, University of Turku, in February 2023, prior to the seasonal antler-shedding period. Antler and metacarpal bone samples were collected immediately after slaughter from 30 reindeer, yielding a total of 60 specimens (30 antlers and 30 metacarpal bones). The sampled animals included 12 males, 15 females, and 3 individuals of unknown sex. Sex was determined on site by experienced reindeer farm personnel based on external morphological characteristics. Sex was recorded as descriptive metadata only and was not included as an experimental factor in the present study. To enable direct within-individual comparisons between antler and metacarpal tissues, only specimens that could be unequivocally matched to the same animal were included in subsequent analyses. Based on this criterion, nine individuals were used for structural, compositional, and histological analyses, whereas eleven individuals were included in proteomic analyses. The bone samples were stored in sterile plastic tubes or bags and frozen at −20 °C until use.

Bone samples of antler tip, antler base, and metacarpus were cut with a diamond saw (Buehler, IL) into slices 1–2 mm in thickness for material characterization and into 10 mm in thickness for proteomics analysis (**Supplementary Fig. S1**). The bone slices were rinsed with ultrasonication in distilled water. The samples were defrosted and cleaned in deionized water for 3 hours at room temperature. The water was exchanged three times, once every hour. After the cleaning step, the samples were allowed to dry thoroughly at room temperature.

### X-ray diffraction (XRD) analysis

The mineral phases of the bone samples were determined with XRD (PANalytical EMPYREAN). The instrument in question uses a Cu X-ray tube which emits Cu Kα radiation with the wavelengths of λKα1 = 1.5406 Å and λKα2 = 1.5444 Å. The scanning was performed at 45kV and 40mA in the 2θ range of 20° to 90°, with a step size of 0.020 degree on a diffraction spectrometer, Bragg-Brentano HD monochromator, and a PIXcel 3D detector.

The resulting diffraction patterns were analyzed in relation to the reference data set recognized by the International Centre for Diffraction Data (ICDD) database. Smoothing and baseline correction were applied to all XRD patterns prior to analysis using Origin software (versions 16, 2023b, and 2025). Parameters including crystallite size, c-axis orientation, and d-spacing were subsequently calculated and compared among reindeer antler tip, antler base, and metacarpal bone samples.

### Fourier-transform infrared spectroscopy (FTIR) analysis

The bone samples were subjected to attenuated total reflectance Fourier transform infrared spectroscopy (ATR-FTIR) using a PerkinElmer Frontier IR with an attached ATR unit. The ATR-FTIR spectra were recorded in the range 400-4000 cm^−1^ with each spectrum representing the average of 10 scans. The transmittance spectra were converted to absorbance with the equation Absorbance = 2 - log(%T), and baseline correction was applied for each spectrum. The parameters such as mineral-to-matrix ratio, mineral maturity, carbonate content/bioapatite, secondary structure of proteins, amide I/phosphate ratio, amide II/phosphate ratio, amide III and collagen/phosphate ratio, amide I and amide II/phosphate ratio, and amides and collagen/phosphate ratio were analyzed using Origin software (version 16, 2023b, and 2025) ^21,22,24^.

### Micro-computed tomography (micro-CT) analysis

Bone samples were scanned using a SkyScan 1272 system (Bruker microCT, Kontich, Belgium) at 45 kV and 40 mA using a 0.5-mm aluminum filter, with a pixel size of 7.5 µm over a 360° rotation. Scanned images were reconstructed with NRecon, version 1.7.5.6 Skyscan, where parameters such as the beam hardening correction, misalignment compensation, ring artifact reduction and the attenuation boundaries were adjusted. The reconstructed results were viewed with DataViewer software as three orthogonal sections. CTAn was used to determine the region of interest (ROI) in each sample.

The reconstructed datasets and ROIs were processed using Bruker’s CT-Analyser (CTAn) and Batman (Batch Manager). Metacarpal cortical bone was included as a mature skeletal reference tissue to facilitate comparisons of microarchitectural and compositional characteristics between regenerating antler tissues and conventional load-bearing bone. Parameters measured for both cortical and trabecular bone were bone mineral density (BMD), total, open, and closed porosity, trabecular number (Tb.N), trabecular separation (Tb.Sp), and trabecular thickness (Tb.Th).

The BMD calibration was performed using Bruker hydroxyapatite phantoms (0.25 and 0.75 g·cm^−3^) scanned under identical conditions. Mean values were calculated from four ROIs within each of the central, intermediate, and peripheral regions of antler tip and antler base samples.

### Histological evaluation

The bone samples (1–2 mm in thickness) prepared as described above were fixed with 4% paraformaldehyde, embedded in paraffin, cut into 5 µm sections, and stained with Masson’s trichrome.

### Statistical analysis for material characterization

To determine whether there were statistically significant differences in the calculated parameters among the three bones, Prism 11 software was used to compute a one-way ANOVA which was followed by Tukey’s Honestly Significant Difference (HSD) test for multiple comparisons.

### Proteomics analysis

#### Sample preparation for proteomics analysis

Bone samples were drilled to acquire bone powder samples. Samples were decalcified with 10 % formic acid for 6 h at +4 °C and centrifuged 14000g for 5 minutes. Supernatant was removed and 6 M guanidine hydrochloride, 100 mM Tris-HCl pH 7.5 -buffer was added to the precipitates and mixed gently. Samples were incubated 18 h at +4 °C. Samples were centrifuged 20000g for 20 minutes and supernatant was collected and digested with FASP ^64^. Samples were added to Microcon YM-10 Centrifugal Filter Units (Merck) and centrifuged for 15 min at 14000g. Proteins were alkylated with 40 mM iodoacetamide for 30 min in the dark at room temperature. Samples were centrifuged for 30 min at 14000g followed by three washes with 50 mM Tris-HCl pH 8.0 with centrifugation for 15 min at 14000g each. Samples were digested overnight with sequencing grade modified trypsin (Promega). After digestion peptide samples were desalted with a Sep-Pak tC18 96- well plate (Waters) and evaporated to dryness. Samples were dissolved in 0.1% formic acid and peptide concentration was determined with a NanoDrop device. For DIA analysis 1000 ng of peptides were injected and analyzed in random order. Wash runs were submitted between each sample to reduce potential carry-over of peptides.

#### LC-ESI-MS/MS Analysis

The LC-ESI-MS/MS analysis was performed on a nanoflow HPLC system (Easy-nLC1200, Thermo Fisher Scientific) coupled to the Orbitrap Lumos Fusion mass spectrometer (Thermo Fisher Scientific, Bremen, Germany) equipped with a nano-electrospray ionization source and FAIMS interface. Peptides were first loaded on a trapping column and subsequently separated inline on a 15 cm C18 column (75 μm x 15 cm, ReproSil-Pur 3 μm 120 Å C18-AQ, Dr. Maisch HPLC GmbH, Ammerbuch-Entringen, Germany). The mobile phase consisted of water with 0.1% formic acid (solvent A) or acetonitrile/water (80:20 (v/v)) with 0.1% formic acid (solvent B). A 120 min gradient was used to elute peptides (60 min from 5% to 21% solvent B followed by 48 min from 21 % to 36 min solvent B and in 6 min from 36 min to 100% of solvent B, followed by 6 min wash stage with solvent B).

Samples were analyzed by a data independent acquisition (DIA) LC-MS/MS method. MS data was acquired automatically by using Thermo Xcalibur 4.6 software (Thermo Fisher Scientific). In a DIA method a duty cycle contained one full scan (400 -1000 m/z) and 30 DIA MS/MS scans covering the mass range 400 -1000 with variable width isolation windows. Data analysis consisted of protein identifications and label-free quantifications of protein abundances. Data was analyzed by Spectronaut software (Biognosys; version 17.6.2). DirectDIA approach was used to identify proteins and label-free quantifications were performed with MaxLFQ.

#### Annotation of reindeer proteins ECM

We used the Phylobone database for the annotation of bone extracellular matrix proteins in reindeer using public data from *Rangifer tarandus* for the annotation and the protocol described in ^65^.

#### Multifactorial analysis

The abundance of proteins was normalized with the housekeeping protein glyceraldehyde-3-phosphate dehydrogenase (GAPDH) by the maximum values within each set of data. The dataset then was log transformed into the final dataset used for analyses (**Tables S1-S7**, **Supplementary dataset 2**). The analysis was performed in spreadsheet and Python v.3.10.

#### Statistical analysis for proteomics analyses

One-tailed Student’s *t*-tests were performed for each protein between antler base, antler tip, and metacarpus pairwise comparisons using a significance threshold of *p* < 0.05. The patterns of significance created eight groups, which were used as the basis for further analysis.

Protein expression within each comparison group was assessed by calculating and comparing the mean abundance of the corresponding combination. A principal component analysis (PCA) was performed to evaluate the structure of the proteomics groups dataset. PCA was conducted with Python’s library Scikit-learn (v.1.8) PCA class. The PCA, box and scattering plots were visualized with the Matplotlib library.

Mesenchymal stem cell isolation, culture, and multilineage differentiation assays were performed as described in the Supplementary Information. Osteogenic, adipogenic, and chondrogenic differentiation was evaluated by Alizarin Red S, Oil Red O, and toluidine blue staining, respectively (**Supplementary Fig. S2**).

## Data availability statements

The authors confirm that the data supporting the findings of this study are available within the article and its Supplementary Information files. Source data underlying the graphs presented in this paper are provided in Supplementary Datasets 1 and 2.

## Acknowledgements

This study was supported by Sigrid Juselius Foundation (#230131), the Japan Society for the Promotion of Science’s Grants-in-Aid for Scientific Research (JP23K08670), the Murata Foundation, the Turku Collegium for Science, Medicine and Technology, and Jenny and Antti Wihuri Foundation. We also thank the continuous collaborative support of the Osteoarthritis Foundation International (OAFI) and the Asociación Española de Pacientes con Osteoporosis (AECOS). This study was supported by the IT-CSC Finland (Project ID #2004931). We thank Erasmus+ undergraduate students Ms. Katja Hrovat for her help in preparing the bone samples and other members and collaborators of the Phylobone project for helpful discussions. We also thank the staff of the Kevo Research Station (especially Jouni Lukkari), the Kaldoaivi Reindeer Owners’ Association (Vesa Länsman), and the team at the Skalluvaara facility (Utsjoki, Lapland, Finland) for their assistance during sample collection. Mass spectrometry analyses were performed at the Turku Proteomics Facility supported by Biocenter Finland.

## Conflict of interest

All authors declare no conflict of interest.

## Contributions

M.N. and P.P. conceived and designed the study. I.S., K.S., M.N., and P.P. collected and processed the reindeer samples. H.S., I.A., P.Pa., M.N. performed XRD analysis. H.S., M.S., T.K., and M.N. performed ATR-FTIR analysis. H.S., S.S., V.J., P.P., and M.N. performed micro-CT analysis. L.T.P. and P.P. performed proteomics analysis. L.T.P. and P.P. implemented bioinformatics pipelines. S.S. and M.N. performed the cell culture experiment. M.N. and P.P. analyzed the data and had full access to the experimental data in the study, and take responsibility for the integrity of the data, and the accuracy of the data analysis. H.S., L.T.P., S.S., M.N., and P.P carried out data acquisition and prepared the manuscript. All authors approved the final version of the manuscript.

## Notes

### Competing Interest Statement

The authors have declared no competing interest.

