## Supplemental information for "Towards Understanding the Microenvironmental Control of Reindeer Antler Regeneration"

### Supplementary methods

### Mesenchymal stem cell (MSC) culture

MSCs were isolated from the bone marrow of reindeer metacarpus (twelve individuals, between 1 and 2 years old, including both males and females), as described in a previous study [1]. Briefly, the metacarpus were dissected from the reindeer immediately after slaughtering. The epiphyses were cut, and the bone marrow was collected into sterilized tubes. The bone marrow cells were maintained in $\alpha$ modified essential medium ($\alpha$MEM; Sigma–Aldrich, MO, USA) supplemented with 10% fetal bovine serum (FBS; Gibco, MA, USA), 100 U/ml penicillin–streptomycin (Gibco, MA, USA) and 20mM N-2-hydroxyethylpiperazine-N-2-ethane sulfonic acid (HEPES; Gibco, MA, USA) in a humidified atmosphere of 5% $CO_2$ in air at 37°C for 2 weeks. After the cells reached 60–70% confluence, they were treated with 0.25% trypsin-EDTA (Gibco, MA, USA) and centrifuged at 1000 rpm for 5 min. The cells were frozen and stored in liquid nitrogen until use. Multilineage differentiation capacity was investigated by differentiating the MSCs into osteoblasts, adipocytes or chondrocytes.

Osteoblasts: The cells were cultured in medium supplemented with $10^{-7}$ M dexamethasone (Sigma–Aldrich, MO, USA), 10 mM β-glycerophosphate (Sigma–Aldrich, MO, USA) and 50 μM ascorbic acid (Sigma–Aldrich, MO, USA) for 6 weeks. The cells were fixed with 4% paraformaldehyde for 20 min and stained with alizarin red S (Sigma–Aldrich, MO, USA).

Adipocytes: The cells were cultured in medium supplemented with $10^{-6}$ M dexamethasone (Sigma–Aldrich, MO, USA), 0.5 μM IBMX (Sigma–Aldrich, MO, USA) and 10 (wt/vol) ng/ml insulin (Sigma–Aldrich, MO, USA) for 6 weeks. The cells were fixed with 4% paraformaldehyde for 20 min and stained with 0.5% oil red O staining solution (Sigma–Aldrich, MO, USA) for 15 min.

Chondrocyte: The cells were cultured in medium supplemented with $10^{-7}$ M dexamethasone (Sigma–Aldrich, MO, USA), 1% ITS (Sigma–Aldrich, MO, USA), 50 μg/ml ascorbic acid (Sigma–Aldrich, MO, USA), 1 mM sodium pyruvate (Sigma–Aldrich, MO, USA), 50 μg/ml (wt/vol) proline (Sigma–Aldrich, MO, USA), or 20 ng/ml (wt/vol) TGF-β3 (Biotechne, MN, USA) for 6 weeks. The cells were fixed with 4% paraformaldehyde for 20 min and stained with 0.5% toluidine blue staining solution (Sigma–Aldrich, MO, USA) for 15 min.

### Supplementary figures

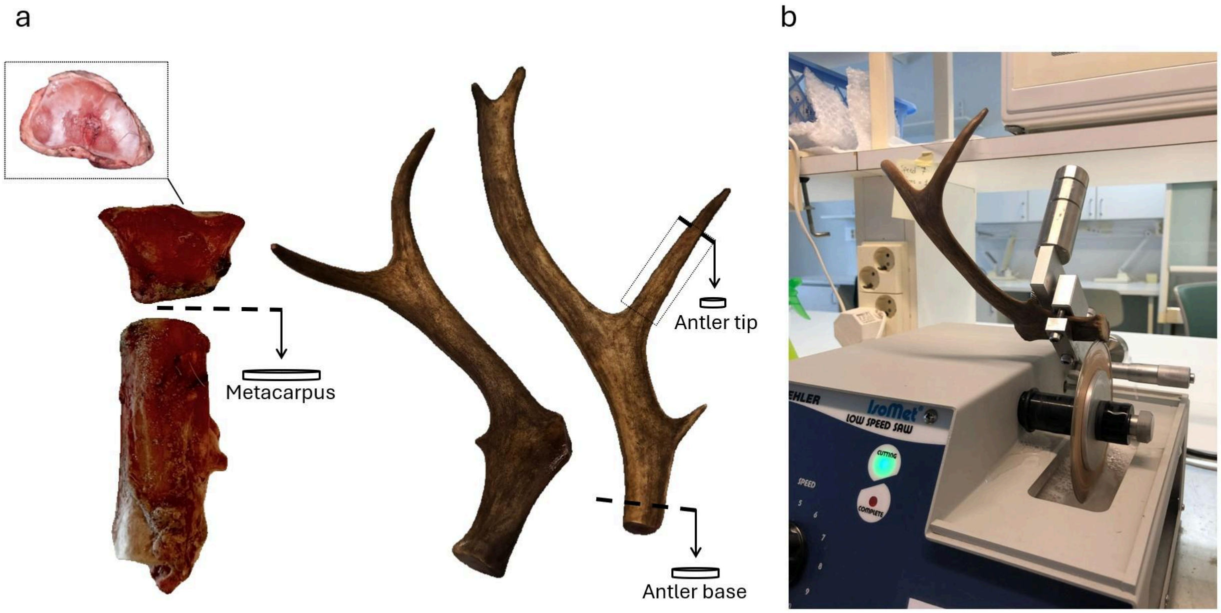


**Fig. S1 Preparation of bone samples. (a)** Bone slices in the thickness of 1–2 mm were obtained from metacarpal diaphysis, antler tips and antler bases. The transverse sample sections of the antler were cut parallel to the vertical axis (dotted lines) while ensuring that the sample surfaces were flat. **(b)** Obtainment of antler slices with a low-speed diamond saw.

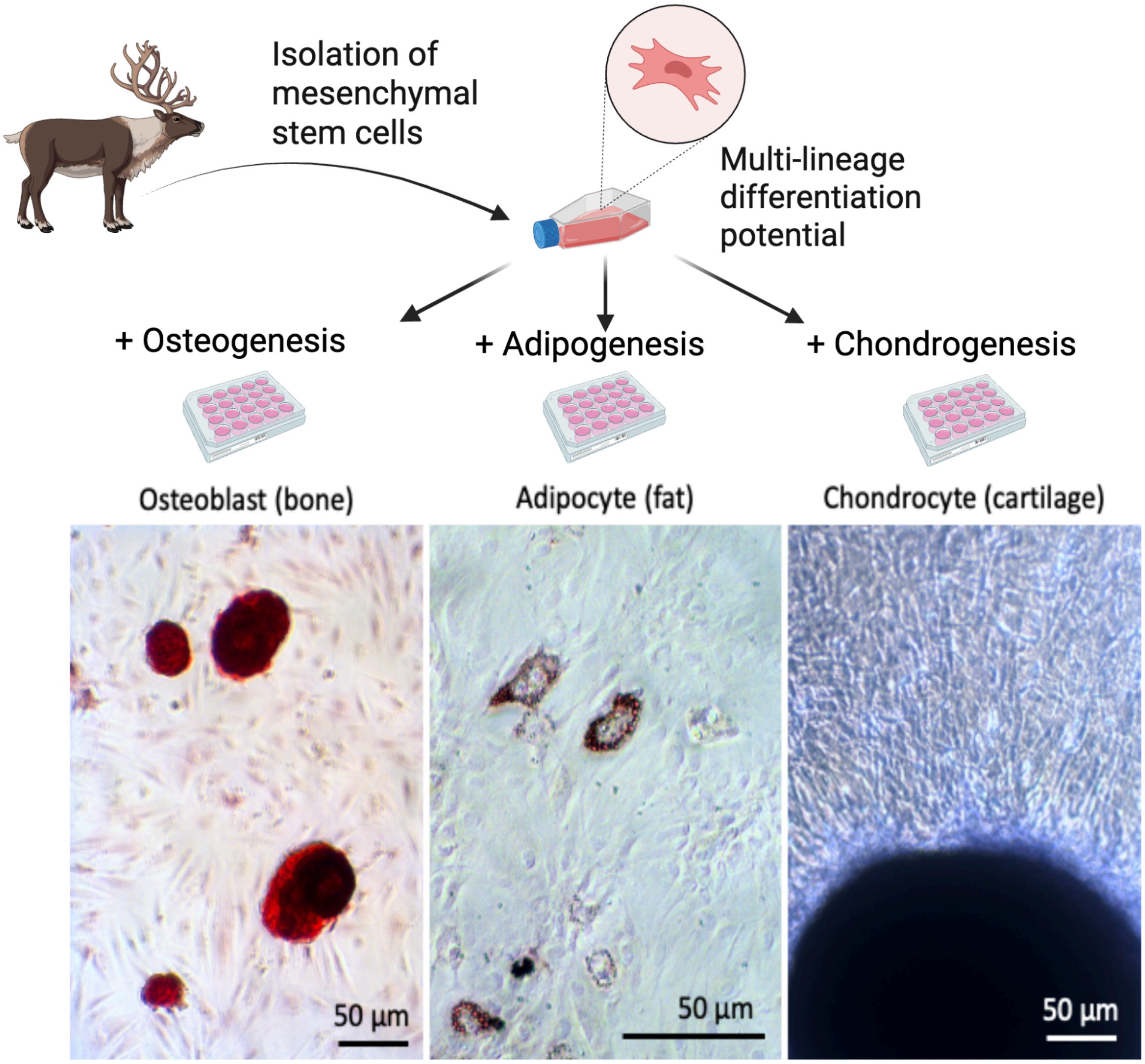


**Fig. S2 Multilineage differentiation of reindeer mesenchymal stem cells.** The images show osteogenic, adipogenic, and chondrogenic differentiation of reindeer mesenchymal stem cells after lineage-specific induction. Osteogenic cultures display intense Alizarin Red S staining of mineralized matrix (left). Adipogenic differentiation is confirmed by the presence of intracellular lipid droplets stained with Oil Red O (middle). Chondrogenic cultures form dense cartilaginous micromass structures with a proteoglycan-rich extracellular matrix (right). Scale bars: 50 µm.

### Supplementary Table

**Table S1.** The log transformed values of 16 proteins that separate metacarpus from tip and base, the respective mean value and expression.

| Samples | MATN4 | CNMD | SFRP5 | CILP | BMP4 | TGFB3 | CCN2 | TNXB | KERA | MATN1 | LAMA3 | TIMP2 | TFPI2 | FST | CHAD | CCL25 |
|---|---|---|---|---|---|---|---|---|---|---|---|---|---|---|---|---|
| Base_1 | -13.51 | -3.76 | -13.51 | -3.80 | -13.51 | -4.19 | -4.21 | -3.81 | -3.59 | -13.51 | -13.51 | -3.86 | -3.86 | -13.51 | -13.51 | -13.51 |
| Base_11 | -8.74 | -4.67 | -8.74 | -8.74 | -8.74 | -5.69 | -8.74 | -5.63 | -8.74 | -8.74 | -8.74 | -4.55 | -8.74 | -8.74 | -8.74 | -8.74 |
| Base_17 | -8.70 | -8.70 | -8.70 | -4.92 | -8.70 | -5.48 | -8.70 | -8.70 | -8.70 | -8.70 | -8.70 | -4.36 | -8.70 | -8.70 | -8.70 | -8.70 |
| Base_20 | -8.54 | -8.54 | -8.54 | -8.54 | -8.54 | -5.61 | -8.54 | -8.54 | -8.54 | -8.54 | -8.54 | -4.13 | -8.54 | -8.54 | -8.54 | -8.54 |
| Base_24 | -8.57 | -8.57 | -8.57 | -4.38 | -5.24 | -8.57 | -8.57 | -5.40 | -8.57 | -8.57 | -8.57 | -4.15 | -8.57 | -8.57 | -8.57 | -8.57 |
| Base_3 | -5.86 | -14.23 | -14.23 | -5.15 | -14.23 | -5.42 | -5.07 | -14.23 | -4.71 | -14.23 | -14.23 | -9.65 | -14.23 | -14.23 | -14.23 | -14.23 |
| Base_4 | -10.61 | -10.61 | -10.61 | -0.91 | -10.61 | -10.61 | -10.61 | -10.61 | -1.47 | -10.61 | -10.61 | -1.08 | -10.61 | -10.61 | -10.61 | -10.61 |
| Base_5 | -13.53 | -3.58 | -13.53 | -3.90 | -13.53 | -13.53 | -13.53 | -4.38 | -4.12 | -13.53 | -13.53 | -3.74 | -13.53 | -13.53 | -13.53 | -13.53 |
| Base_6 | -8.88 | -8.88 | -8.88 | -8.88 | -8.88 | -8.88 | -8.88 | -8.88 | -4.88 | -8.88 | -8.88 | -4.32 | -8.88 | -8.88 | -8.88 | -8.88 |
| Base_7 | -9.11 | -9.11 | -9.11 | -9.11 | -9.11 | -9.11 | -9.11 | -6.20 | -5.71 | -9.11 | -9.11 | -4.88 | -9.11 | -9.11 | -9.11 | -9.11 |
| Base_8 | -8.53 | -4.49 | -8.53 | -8.53 | -8.53 | -8.53 | -8.53 | -8.53 | -8.53 | -8.53 | -8.53 | -3.68 | -8.53 | -8.53 | -8.53 | -8.53 |
| Metacarpus_1 | -3.92 | -4.00 | -13.42 | -4.39 | -3.69 | -3.82 | -3.60 | -4.35 | -7.69 | -5.32 | -3.50 | -3.60 | -4.03 | -4.40 | -13.42 | -3.90 |
| Metacarpus_11 | -8.53 | -3.88 | -8.53 | -4.24 | -4.81 | -4.18 | -8.53 | -5.61 | -3.26 | -8.53 | -8.53 | -3.29 | -8.53 | -5.61 | -8.53 | -8.53 |
| Metacarpus_17 | -8.57 | -4.09 | -8.57 | -4.34 | -5.22 | -4.86 | -8.57 | -5.11 | -4.24 | -8.57 | -8.57 | -3.41 | -8.57 | -5.09 | -8.57 | -8.57 |
| Metacarpus_20 | -8.53 | -3.97 | -8.53 | -4.21 | -5.60 | -4.51 | -8.53 | -4.96 | -3.13 | -8.53 | -8.53 | -3.25 | -8.53 | -8.53 | -8.53 | -8.53 |
| Metacarpus_24 | -8.56 | -4.11 | -8.56 | -8.56 | -4.84 | -5.06 | -8.56 | -5.39 | -4.81 | -8.56 | -8.56 | -3.49 | -8.56 | -4.36 | -8.56 | -8.56 |
| Metacarpus_3 | -4.24 | -4.30 | -3.90 | -4.02 | -3.80 | -4.23 | -4.46 | -4.44 | -3.57 | -3.59 | -4.09 | -4.03 | -3.88 | -3.65 | -3.56 | -4.36 |
| Metacarpus_4 | -4.20 | -4.14 | -3.83 | -4.14 | -4.18 | -5.42 | -4.03 | -3.76 | -4.19 | -4.00 | -4.16 | -3.53 | -3.67 | -13.51 | -3.75 | -3.82 |
| Metacarpus_5 | -1.22 | -0.81 | -0.78 | -0.55 | -1.30 | -0.71 | -1.18 | -1.18 | -0.94 | -0.67 | -0.98 | -1.31 | -0.68 | -0.73 | -0.49 | -0.46 |
| Metacarpus_6 | -8.36 | -8.36 | -8.36 | -3.82 | -4.30 | -3.43 | -8.36 | -8.36 | -3.06 | -8.36 | -8.36 | -2.56 | -8.36 | -4.36 | -8.36 | -8.36 |
| Metacarpus_7 | -8.48 | -8.48 | -8.48 | -3.82 | -4.84 | -4.71 | -8.48 | -5.02 | -3.28 | -8.48 | -8.48 | -3.19 | -8.48 | -8.48 | -8.48 | -8.48 |
| Metacarpus_8 | -8.58 | -4.15 | -8.58 | -4.48 | -4.50 | -5.12 | -8.58 | -5.18 | -3.83 | -8.58 | -8.58 | -3.39 | -8.58 | -5.31 | -8.58 | -8.58 |
| Tip_1 | -13.46 | -13.46 | -13.46 | -4.38 | -13.46 | -13.46 | -13.46 | -3.65 | -3.84 | -13.46 | -13.46 | -13.46 | -13.46 | -13.46 | -13.46 | -13.46 |
| Tip_11 | -8.30 | -8.30 | -8.30 | -4.56 | -8.30 | -5.11 | -8.30 | -5.77 | -8.30 | -8.30 | -8.30 | -4.17 | -8.30 | -8.30 | -8.30 | -8.30 |
| Tip_17 | -8.48 | -8.48 | -8.48 | -3.85 | -8.48 | -8.48 | -8.48 | -8.48 | -8.48 | -8.48 | -8.48 | -4.58 | -8.48 | -8.48 | -8.48 | -8.48 |
| Tip_20 | -8.65 | -8.65 | -8.65 | -5.14 | -8.65 | -5.44 | -8.65 | -5.74 | -5.54 | -8.65 | -8.65 | -4.69 | -8.65 | -8.65 | -8.65 | -8.65 |
| Tip_24 | -7.78 | -7.78 | -7.78 | -7.78 | -3.66 | -7.78 | -7.78 | -7.78 | -7.78 | -7.78 | -7.78 | -3.13 | -7.78 | -7.78 | -7.78 | -7.78 |
| Tip_3 | -9.99 | -0.81 | -9.99 | -9.99 | -0.07 | -0.77 | -9.99 | -9.99 | -0.56 | -9.99 | -9.99 | -1.79 | -9.99 | -0.38 | -9.99 | -9.99 |
| Tip_4 | -4.12 | -5.04 | -9.60 | -3.53 | -3.44 | -4.00 | -13.30 | -3.39 | -4.80 | -13.30 | -13.30 | -4.17 | -3.37 | -13.30 | -13.30 | -3.42 |
| Tip_5 | -13.92 | -13.92 | -13.92 | -4.52 | -13.92 | -4.71 | -13.92 | -13.92 | -4.52 | -13.92 | -4.56 | -4.85 | -13.92 | -13.92 | -13.92 | -13.92 |
| Tip_6 | -8.55 | -8.55 | -8.55 | -8.55 | -8.55 | -8.55 | -8.55 | -5.59 | -8.55 | -8.55 | -8.55 | -4.47 | -8.55 | -8.55 | -8.55 | -8.55 |
| Tip_7 | -8.47 | -4.74 | -8.47 | -8.47 | -8.47 | -8.47 | -8.47 | -5.81 | -8.47 | -8.47 | -8.47 | -4.27 | -8.47 | -8.47 | -8.47 | -8.47 |
| Tip_8 | -7.99 | -7.99 | -7.99 | -7.99 | -4.11 | -7.99 | -7.99 | -7.99 | -7.99 | -7.99 | -7.99 | -3.19 | -7.99 | -3.81 | -7.99 | -7.99 |
| **Mean Base+Tip** | **-9.29** | **-7.86** | **-9.91** | **-6.16** | **-8.67** | **-7.29** | **-9.24** | **-7.41** | **-6.20** | **-10.08** | **-9.66** | **-4.60** | **-9.19** | **-9.46** | **-10.08** | **-9.63** |
| **Mean Metacarpus** | **-6.65** | **-4.57** | **-7.41** | **-4.23** | **-4.28** | **-4.19** | **-6.63** | **-4.85** | **-3.82** | **-6.65** | **-6.58** | **-3.19** | **-6.53** | **-5.82** | **-7.35** | **-6.56** |
| **Base+Tip** | - | - | - | - | - | - | - | - | - | - | - | - | - | - | - | - |
| **Metacarpus** | + | + | + | + | + | + | + | + | + | + | + | + | + | + | + | + |

**Table S2.** The log transformed values of 4 proteins that separate tip from metacarpus and base, the respective mean value and expression.

| Samples | ITIH6 | COL8A1 | COL16A1 | MSTN |
|---|---|---|---|---|
| Base_1 | -3.74 | -3.85 | -4.31 | -4.06 |
| Base_11 | -5.61 | -5.32 | -4.80 | -5.39 |
| Base_17 | -5.49 | -4.85 | -4.51 | -4.66 |
| Base_20 | -5.28 | -4.93 | -3.86 | -8.54 |
| Base_24 | -7.17 | -5.26 | -3.87 | -4.90 |
| Base_3 | -4.39 | -4.87 | -5.14 | -4.35 |
| Base_4 | -1.29 | -0.95 | -1.20 | -1.54 |
| Base_5 | -4.44 | -4.41 | -4.45 | -4.35 |
| Base_6 | -5.22 | -5.20 | -6.26 | -5.71 |
| Base_7 | -5.13 | -9.11 | -4.96 | -9.11 |
| Base_8 | -4.90 | -4.74 | -8.53 | -5.55 |
| Metacarpus_1 | -4.36 | -4.19 | -3.70 | -4.11 |
| Metacarpus_11 | -5.11 | -4.34 | -4.24 | -4.62 |
| Metacarpus_17 | -5.29 | -4.75 | -8.57 | -4.80 |
| Metacarpus_20 | -5.69 | -3.97 | -8.53 | -4.82 |
| Metacarpus_24 | -5.16 | -3.96 | -4.10 | -5.15 |
| Metacarpus_3 | -3.63 | -3.60 | -3.58 | -3.56 |
| Metacarpus_4 | -3.89 | -8.36 | -4.98 | -3.66 |
| Metacarpus_5 | -1.35 | -0.60 | -1.38 | -1.91 |
| Metacarpus_6 | -4.56 | -4.57 | -3.43 | -4.11 |
| Metacarpus_7 | -4.90 | -3.78 | -3.85 | -4.66 |
| Metacarpus_8 | -5.22 | -4.70 | -8.58 | -4.82 |
| Tip_1 | -13.46 | -13.46 | -3.79 | -13.46 |
| Tip_11 | -5.41 | -5.12 | -3.57 | -8.30 |
| Tip_17 | -4.38 | -5.30 | -3.91 | -8.48 |
| Tip_20 | -6.31 | -5.69 | -4.38 | -8.65 |
| Tip_24 | -7.78 | -7.78 | -2.52 | -7.78 |
| Tip_3 | -9.99 | -9.99 | -0.41 | -9.99 |
| Tip_4 | -4.65 | -4.59 | -3.78 | -4.23 |
| Tip_5 | -13.92 | -4.80 | -4.20 | -4.01 |
| Tip_6 | -4.96 | -5.18 | -3.74 | -8.55 |
| Tip_7 | -5.32 | -5.12 | -3.86 | -8.47 |
| Tip_8 | -7.99 | -7.99 | -2.93 | -7.99 |
| **Mean Base+Metacarpus** | **-4.63** | **-4.56** | **-4.86** | **-4.74** |
| **Mean Tip** | **-7.65** | **-6.82** | **-3.37** | **-8.17** |
| **Base+Metacarpus** | + | + | - | + |
| **Tip** | - | - | + | - |

**Table S3.** The log transformed values of 10 proteins that separate base from tip and metacarpus, the respective mean value and expression.

| Samples | MELTF | FNDC1 | LRRC17\|TNC | LYZ | NDNF | ACAN | ESM1 | SH3PXD2B | HAPLN1 | COL5A3 |
|---|---|---|---|---|---|---|---|---|---|---|
| Base_1 | -3.60 | -4.11 | -3.77 | -3.72 | -13.51 | -4.17 | -13.51 | -3.70 | -3.75 | -4.47 |
| Base_11 | -5.45 | -5.22 | -3.93 | -8.74 | -8.74 | -5.91 | -8.74 | -4.57 | -8.74 | -4.53 |
| Base_17 | -5.03 | -4.81 | -3.85 | -5.11 | -8.70 | -5.01 | -8.70 | -4.44 | -8.70 | -4.33 |
| Base_20 | -5.26 | -5.56 | -3.24 | -4.80 | -8.54 | -4.59 | -8.54 | -4.09 | -5.24 | -4.43 |
| Base_24 | -7.54 | -5.07 | -3.74 | -4.87 | -8.57 | -4.95 | -8.57 | -4.39 | -8.57 | -4.07 |
| Base_3 | -14.23 | -6.08 | -5.11 | -14.23 | -5.49 | -4.41 | -14.23 | -6.21 | -4.29 | -4.74 |
| Base_4 | -10.61 | -1.15 | -5.85 | -0.62 | -3.38 | -6.43 | -10.61 | -10.61 | -1.27 | -10.61 |
| Base_5 | -3.72 | -4.07 | -4.47 | -13.53 | -4.74 | -3.93 | -13.53 | -3.61 | -4.74 | -13.53 |
| Base_6 | -5.52 | -5.24 | -4.50 | -3.46 | -8.88 | -4.99 | -8.88 | -4.97 | -8.88 | -4.79 |
| Base_7 | -9.11 | -6.16 | -4.32 | -9.11 | -9.11 | -5.60 | -9.11 | -5.15 | -9.11 | -4.82 |
| Base_8 | -4.51 | -4.72 | -3.58 | -4.73 | -8.53 | -4.39 | -8.53 | -4.15 | -8.53 | -4.00 |
| Metacarpus_1 | -4.11 | -4.31 | -3.95 | -3.52 | -4.24 | -4.15 | -3.65 | -4.31 | -4.41 | -3.96 |
| Metacarpus_11 | -4.24 | -3.81 | -3.37 | -3.63 | -4.93 | -3.85 | -8.53 | -4.19 | -4.56 | -4.14 |
| Metacarpus_17 | -4.30 | -3.95 | -3.04 | -4.05 | -4.70 | -4.08 | -8.57 | -4.27 | -4.82 | -4.21 |
| Metacarpus_20 | -4.27 | -3.80 | -2.81 | -3.92 | -5.38 | -3.23 | -8.53 | -4.19 | -4.57 | -4.16 |
| Metacarpus_24 | -3.64 | -4.02 | -4.03 | -3.91 | -5.10 | -4.41 | -8.56 | -4.34 | -4.28 | -4.39 |
| Metacarpus_3 | -3.99 | -4.42 | -4.06 | -3.69 | -3.99 | -3.53 | -3.76 | -3.75 | -4.46 | -5.92 |
| Metacarpus_4 | -3.98 | -4.41 | -4.14 | -4.33 | -4.01 | -4.22 | -3.89 | -3.93 | -3.68 | -4.36 |
| Metacarpus_5 | -1.28 | -0.45 | -1.04 | -0.62 | -2.17 | -1.33 | -0.70 | -0.96 | -0.52 | -0.45 |
| Metacarpus_6 | -3.52 | -3.46 | -2.90 | -3.17 | -8.36 | -3.23 | -8.36 | -3.91 | -3.90 | -3.60 |
| Metacarpus_7 | -3.70 | -3.62 | -2.66 | -3.10 | -4.53 | -3.48 | -8.48 | -4.16 | -4.27 | -3.42 |
| Metacarpus_8 | -4.29 | -3.82 | -3.49 | -3.81 | -5.07 | -3.87 | -8.58 | -4.32 | -5.05 | -4.02 |
| Tip_1 | -3.71 | -4.16 | -3.90 | -3.61 | -5.79 | -4.41 | -5.16 | -5.36 | -3.54 | -4.45 |
| Tip_11 | -4.64 | -4.10 | -3.29 | -4.91 | -5.06 | -4.12 | -8.30 | -3.89 | -4.85 | -3.91 |
| Tip_17 | -4.99 | -4.48 | -3.52 | -4.50 | -8.48 | -4.63 | -8.48 | -4.13 | -8.48 | -3.95 |
| Tip_20 | -6.14 | -4.41 | -3.35 | -5.01 | -5.53 | -4.00 | -8.65 | -3.89 | -4.96 | -4.66 |
| Tip_24 | -3.46 | -2.96 | -2.07 | -3.99 | -3.86 | -3.45 | -7.78 | -3.63 | -3.81 | -2.63 |
| Tip_3 | -0.51 | -0.87 | -3.79 | -0.58 | -0.48 | -4.84 | -9.99 | -0.86 | -0.91 | -0.81 |
| Tip_4 | -3.64 | -3.42 | -3.33 | -3.46 | -3.85 | -3.34 | -3.75 | -4.40 | -3.55 | -4.20 |
| Tip_5 | -3.99 | -4.48 | -4.44 | -4.10 | -4.14 | -4.33 | -13.92 | -4.39 | -5.58 | -3.96 |
| Tip_6 | -4.88 | -4.29 | -3.43 | -3.98 | -4.85 | -4.40 | -8.55 | -4.25 | -5.19 | -3.52 |
| Tip_7 | -5.07 | -4.28 | -3.37 | -5.07 | -5.24 | -4.07 | -8.47 | -3.88 | -4.89 | -4.17 |
| Tip_8 | -3.54 | -3.32 | -2.39 | -3.59 | -7.99 | -3.33 | -7.99 | -3.28 | -4.32 | -2.87 |
| **Mean Metacarpus+Tip** | **-3.90** | **-3.67** | **-3.29** | **-3.66** | **-4.90** | **-3.83** | **-7.39** | **-3.83** | **-4.30** | **-3.72** |
| **Mean Base** | **-6.78** | **-4.75** | **-4.21** | **-6.63** | **-8.02** | **-4.94** | **-10.27** | **-5.08** | **-6.53** | **-5.85** |
| **Tip+Metacarpus** | + | + | + | + | + | + | + | + | + | + |
| **Base** | - | - | - | - | - | - | - | - | - | - |

**Table S4.** The log transformed values of 13 proteins that separate metacarpus and base, the respective mean value and expression.

| Samples | TNN | TGFB2 | COL9A3\|COL9A2 | HBA1 | THBS2 | BMP3 | ENPP1 | TINAGL1 | BMPER | MFGE8 | CTSG | TCN2 | SFRP1 |
|---|---|---|---|---|---|---|---|---|---|---|---|---|---|
| Base_1 | -4.00 | -8.94 | -13.51 | -4.51 | -3.53 | -4.00 | -3.65 | -4.24 | -3.96 | -3.62 | -4.07 | -4.45 | -3.52 |
| Base_11 | -4.66 | -4.93 | -8.74 | -4.31 | -4.30 | -4.59 | -5.26 | -4.54 | -4.62 | -3.59 | -8.74 | -3.90 | -4.56 |
| Base_17 | -4.78 | -4.38 | -8.70 | -4.41 | -4.36 | -4.44 | -4.87 | -4.89 | -4.74 | -3.41 | -8.70 | -3.77 | -5.46 |
| Base_20 | -4.03 | -4.71 | -8.54 | -3.99 | -4.16 | -4.36 | -8.54 | -4.71 | -4.44 | -3.31 | -4.65 | -3.79 | -4.43 |
| Base_24 | -4.42 | -4.52 | -8.57 | -4.51 | -3.78 | -3.97 | -4.66 | -4.87 | -4.22 | -3.28 | -6.84 | -3.49 | -5.51 |
| Base_3 | -4.28 | -5.00 | -14.23 | -4.23 | -4.91 | -5.18 | -4.24 | -4.94 | -4.80 | -5.08 | -5.20 | -4.84 | -4.77 |
| Base_4 | -1.31 | -1.30 | -10.61 | -1.09 | -1.24 | -1.59 | -1.37 | -2.19 | -10.61 | -1.41 | -0.93 | -5.23 | -2.21 |
| Base_5 | -3.60 | -4.22 | -13.53 | -4.38 | -4.38 | -3.61 | -4.31 | -5.45 | -13.53 | -5.38 | -3.67 | -4.25 | -13.53 |
| Base_6 | -4.36 | -4.79 | -8.88 | -4.99 | -4.56 | -4.24 | -5.10 | -4.11 | -8.88 | -3.90 | -8.88 | -4.19 | -8.88 |
| Base_7 | -4.93 | -5.63 | -9.11 | -4.22 | -4.89 | -5.07 | -9.11 | -4.62 | -9.11 | -4.21 | -4.82 | -4.34 | -5.24 |
| Base_8 | -4.82 | -4.28 | -8.53 | -4.42 | -4.18 | -4.17 | -4.92 | -4.85 | -4.35 | -3.01 | -8.53 | -3.53 | -4.49 |
| Metacarpus_1 | -3.49 | -4.79 | -13.42 | -4.14 | -3.63 | -4.64 | -3.64 | -4.37 | -3.90 | -3.70 | -13.42 | -3.75 | -4.38 |
| Metacarpus_11 | -3.25 | -3.62 | -8.53 | -8.53 | -3.36 | -2.99 | -4.04 | -4.13 | -3.48 | -2.44 | -8.53 | -3.07 | -3.66 |
| Metacarpus_17 | -2.97 | -3.80 | -8.57 | -8.57 | -3.50 | -3.38 | -4.38 | -4.21 | -3.40 | -2.60 | -8.57 | -3.41 | -3.85 |
| Metacarpus_20 | -2.80 | -3.80 | -8.53 | -8.53 | -3.70 | -3.35 | -4.26 | -4.35 | -3.27 | -2.59 | -8.53 | -3.20 | -3.89 |
| Metacarpus_24 | -3.66 | -3.85 | -8.56 | -8.56 | -3.29 | -3.45 | -4.33 | -3.68 | -8.56 | -2.66 | -8.56 | -3.24 | -4.84 |
| Metacarpus_3 | -4.43 | -4.66 | -4.27 | -3.79 | -3.96 | -3.76 | -4.16 | -4.35 | -4.36 | -4.20 | -4.32 | -4.38 | -3.90 |
| Metacarpus_4 | -3.90 | -3.57 | -3.69 | -13.51 | -3.63 | -3.63 | -3.73 | -4.21 | -3.95 | -3.78 | -13.51 | -3.71 | -3.75 |
| Metacarpus_5 | -1.04 | -0.71 | -10.42 | -1.00 | -0.59 | -1.90 | -0.76 | -0.57 | -0.85 | -0.75 | -0.93 | -0.81 | -0.51 |
| Metacarpus_6 | -2.82 | -3.49 | -8.36 | -4.47 | -3.30 | -3.07 | -3.45 | -3.59 | -3.04 | -2.35 | -8.36 | -3.03 | -3.95 |
| Metacarpus_7 | -3.23 | -3.63 | -8.48 | -8.48 | -3.35 | -3.18 | -4.36 | -3.72 | -3.02 | -2.68 | -8.48 | -3.04 | -3.63 |
| Metacarpus_8 | -3.18 | -3.79 | -8.58 | -6.09 | -3.52 | -3.53 | -4.42 | -4.00 | -3.72 | -2.71 | -8.58 | -3.32 | -3.89 |
| Tip_1 | -4.33 | -4.11 | -13.46 | -13.46 | -4.40 | -4.36 | -3.87 | -3.55 | -3.57 | -4.06 | -13.46 | -4.17 | -3.56 |
| Tip_11 | -3.72 | -4.48 | -8.30 | -4.56 | -3.15 | -4.76 | -4.87 | -5.07 | -4.47 | -3.32 | -4.95 | -3.81 | -4.91 |
| Tip_17 | -4.19 | -4.54 | -8.48 | -4.00 | -3.67 | -4.67 | -5.26 | -4.72 | -4.14 | -3.43 | -8.48 | -3.77 | -5.18 |
| Tip_20 | -3.70 | -5.45 | -8.65 | -5.36 | -3.76 | -5.45 | -5.68 | -5.05 | -8.65 | -3.99 | -5.28 | -4.56 | -5.21 |
| Tip_24 | -2.56 | -3.62 | -7.78 | -7.78 | -2.02 | -3.61 | -3.90 | -4.04 | -7.78 | -1.86 | -7.78 | -2.42 | -3.72 |
| Tip_3 | -1.42 | -0.67 | -9.99 | -0.13 | -3.74 | -0.09 | -2.14 | -0.40 | -0.34 | -0.85 | -0.24 | -0.86 | -1.77 |
| Tip_4 | -4.00 | -3.98 | -4.03 | -4.04 | -3.32 | -3.87 | -3.88 | -3.86 | -3.71 | -4.29 | -4.22 | -4.20 | -4.08 |
| Tip_5 | -4.80 | -3.97 | -13.92 | -4.51 | -4.11 | -4.32 | -3.95 | -4.35 | -4.65 | -8.93 | -4.48 | -4.92 | -4.72 |
| Tip_6 | -3.96 | -4.93 | -8.55 | -4.76 | -3.26 | -4.35 | -5.01 | -4.74 | -4.39 | -3.35 | -8.55 | -3.96 | -4.53 |
| Tip_7 | -3.61 | -4.57 | -8.47 | -5.15 | -3.34 | -4.80 | -5.25 | -5.16 | -8.47 | -3.39 | -8.47 | -3.97 | -4.69 |
| Tip_8 | -2.69 | -3.63 | -7.99 | -7.99 | -2.94 | -4.01 | -3.77 | -4.07 | -7.99 | -2.06 | -7.99 | -2.54 | -3.78 |
| **Mean Metacarpus** | **-3.16** | **-3.61** | **-8.31** | **-6.88** | **-3.26** | **-3.35** | **-3.78** | **-3.74** | **-3.78** | **-2.77** | **-8.35** | **-3.18** | **-3.66** |
| **Mean Base** | **-4.11** | **-4.79** | **-10.27** | **-4.10** | **-4.03** | **-4.11** | **-5.09** | **-4.49** | **-6.66** | **-3.66** | **-5.91** | **-4.16** | **-5.69** |
| **Metacarpus** | + | + | + | - | + | + | + | + | + | + | - | + | + |
| **Base** | - | - | - | + | - | - | - | - | - | - | + | - | - |

**Table 5.** The log transformed values of 2 proteins that separate tip and base, the respective mean value and expression.

| Samples | SERPINB5 | VCAN |
|---|---|---|
| Base_1 | -13.51 | -4.66 |
| Base_11 | -5.35 | -3.71 |
| Base_17 | -8.70 | -3.53 |
| Base_20 | -4.23 | -3.68 |
| Base_24 | -8.57 | -3.40 |
| Base_3 | -14.23 | -5.15 |
| Base_4 | -10.61 | -5.34 |
| Base_5 | -5.46 | -4.36 |
| Base_6 | -3.64 | -3.62 |
| Base_7 | -9.11 | -4.24 |
| Base_8 | -8.53 | -3.02 |
| Metacarpus_1 | -3.66 | -7.16 |
| Metacarpus_11 | -6.11 | -2.49 |
| Metacarpus_17 | -8.57 | -2.45 |
| Metacarpus_20 | -8.53 | -2.48 |
| Metacarpus_24 | -8.56 | -2.27 |
| Metacarpus_3 | -13.51 | -3.92 |
| Metacarpus_4 | -3.71 | -7.32 |
| Metacarpus_5 | -0.72 | -1.20 |
| Metacarpus_6 | -8.36 | -2.15 |
| Metacarpus_7 | -8.48 | -2.16 |
| Metacarpus_8 | -8.58 | -2.39 |
| Tip_1 | -13.46 | -4.10 |
| Tip_11 | -3.88 | -3.56 |
| Tip_17 | -8.48 | -3.60 |
| Tip_20 | -4.30 | -3.75 |
| Tip_24 | -7.78 | -2.33 |
| Tip_3 | -0.08 | -0.78 |
| Tip_4 | -3.41 | -3.32 |
| Tip_5 | -4.03 | -3.97 |
| Tip_6 | -4.78 | -3.61 |
| Tip_7 | -4.15 | -3.59 |
| Tip_8 | -7.99 | -2.41 |
| **Mean Tip** | **-5.67** | **-3.18** |
| **Mean Base** | **-8.36** | **-4.07** |
| **Tip** | + | + |
| **Base** | - | - |

**Table S6.** The log transformed values of 7 proteins that separate metacarpus and tip, the respective mean value and expression.

| Samples | ADPGK | PON2 (CAI9699503.1) | PON2 (CAI9699501.1) | LAMB4\|LAMB1 | ZYX | FRZB | APOA4 |
|---|---|---|---|---|---|---|---|
| Base_1 | -4.40 | -4.89 | -4.25 | -3.91 | -4.16 | -4.39 | -7.18 |
| Base_11 | -5.02 | -4.55 | -6.44 | -5.02 | -8.74 | -4.94 | -2.88 |
| Base_17 | -4.66 | -4.14 | -5.11 | -4.79 | -8.70 | -8.70 | -2.83 |
| Base_20 | -5.03 | -5.01 | -5.96 | -4.79 | -8.54 | -4.57 | -3.24 |
| Base_24 | -4.97 | -3.77 | -6.24 | -4.93 | -8.57 | -8.57 | -2.41 |
| Base_3 | -4.44 | -4.62 | -4.89 | -4.68 | -14.23 | -5.09 | -5.21 |
| Base_4 | -1.59 | -1.54 | -1.12 | -1.99 | -0.88 | -1.12 | -1.29 |
| Base_5 | -3.74 | -4.26 | -4.53 | -4.07 | -4.25 | -13.53 | -7.40 |
| Base_6 | -4.59 | -4.57 | -4.80 | -4.79 | -8.88 | -5.09 | -3.21 |
| Base_7 | -5.23 | -4.89 | -6.88 | -4.89 | -9.11 | -9.11 | -3.37 |
| Base_8 | -4.97 | -3.82 | -4.08 | -4.52 | -8.53 | -8.53 | -2.72 |
| Metacarpus_1 | -4.60 | -13.42 | -13.42 | -4.20 | -4.65 | -4.03 | -7.55 |
| Metacarpus_11 | -5.50 | -5.34 | -7.17 | -5.78 | -8.53 | -4.47 | -2.98 |
| Metacarpus_17 | -5.31 | -5.11 | -6.32 | -4.95 | -8.57 | -4.60 | -3.19 |
| Metacarpus_20 | -5.54 | -5.02 | -6.21 | -4.58 | -8.53 | -4.37 | -3.42 |
| Metacarpus_24 | -5.03 | -4.71 | -6.30 | -8.56 | -8.56 | -8.56 | -2.97 |
| Metacarpus_3 | -4.92 | -4.53 | -3.70 | -4.22 | -3.77 | -3.97 | -4.14 |
| Metacarpus_4 | -4.42 | -3.97 | -13.51 | -6.45 | -3.98 | -8.47 | -4.61 |
| Metacarpus_5 | -1.32 | -1.25 | -0.72 | -1.16 | -0.76 | -0.82 | -4.60 |
| Metacarpus_6 | -4.54 | -8.36 | -5.99 | -4.53 | -8.36 | -4.01 | -2.91 |
| Metacarpus_7 | -5.76 | -4.66 | -5.90 | -5.41 | -8.48 | -8.48 | -3.20 |
| Metacarpus_8 | -4.77 | -5.94 | -6.73 | -4.96 | -8.58 | -8.58 | -3.32 |
| Tip_1 | -4.41 | -3.80 | -3.70 | -4.39 | -13.46 | -13.46 | -4.40 |
| Tip_11 | -4.15 | -4.70 | -4.98 | -4.22 | -8.30 | -8.30 | -3.31 |
| Tip_17 | -4.41 | -4.16 | -5.07 | -4.27 | -8.48 | -8.48 | -2.73 |
| Tip_20 | -4.60 | -5.46 | -5.07 | -4.33 | -8.65 | -8.65 | -3.73 |
| Tip_24 | -3.47 | -3.01 | -4.87 | -3.62 | -7.78 | -7.78 | -1.60 |
| Tip_3 | -1.97 | -0.39 | -0.17 | -0.98 | -9.99 | -9.99 | -1.21 |
| Tip_4 | -3.58 | -3.86 | -3.68 | -3.82 | -3.33 | -4.14 | -3.38 |
| Tip_5 | -4.12 | -4.79 | -5.04 | -4.39 | -13.92 | -4.46 | -4.30 |
| Tip_6 | -4.60 | -3.84 | -5.71 | -4.19 | -8.55 | -5.23 | -2.76 |
| Tip_7 | -4.02 | -4.62 | -4.38 | -4.27 | -8.47 | -5.44 | -3.35 |
| Tip_8 | -3.62 | -3.36 | -4.59 | -3.98 | -7.99 | -7.99 | -1.94 |
| **Mean Metacarpus** | **-4.70** | **-5.66** | **-6.91** | **-4.98** | **-6.62** | **-5.49** | **-3.90** |
| **Mean Tip** | **-3.91** | **-3.82** | **-4.30** | **-3.86** | **-8.99** | **-7.63** | **-2.97** |
| **Metacarpus** | - | - | - | - | + | + | - |
| **Tip** | + | + | + | + | - | - | + |

**Table S7.** The log transformed values of 1 protein that separate metacarpus, tip and base from each other, the respective mean value and expression.

| Samples | PRTN3 |
|---|---|
| Base_1 | -7.49 |
| Base_11 | -3.21 |
| Base_17 | -3.69 |
| Base_20 | -4.37 |
| Base_24 | -3.45 |
| Base_3 | -4.36 |
| Base_4 | -0.74 |
| Base_5 | -3.66 |
| Base_6 | -2.69 |
| Base_7 | -3.04 |
| Base_8 | -8.53 |
| Metacarpus_1 | -13.42 |
| Metacarpus_11 | -8.53 |
| Metacarpus_17 | -8.57 |
| Metacarpus_20 | -8.53 |
| Metacarpus_24 | -8.56 |
| Metacarpus_3 | -13.51 |
| Metacarpus_4 | -13.51 |
| Metacarpus_5 | -10.42 |
| Metacarpus_6 | -8.36 |
| Metacarpus_7 | -8.48 |
| Metacarpus_8 | -8.58 |
| Tip_1 | -4.35 |
| Tip_11 | -4.63 |
| Tip_17 | -8.48 |
| Tip_20 | -5.30 |
| Tip_24 | -7.78 |
| Tip_3 | -9.99 |
| Tip_4 | -3.94 |
| Tip_5 | -3.98 |
| Tip_6 | -5.12 |
| Tip_7 | -8.47 |
| Tip_8 | -7.99 |
| **Mean Metacarpus** | **-10.04** |
| **Mean Base** | **-4.11** |
| **Mean Tip** | **-6.37** |
| **Metacarpus** | - |
| **Base** | + |

**Table S8.** Putative role of bone/antler extracellular matrix (ECM) proteins on the regulation of osteoblast and osteoclast.

| Protein | MC | Antler Base | Antler Tip | Positive Association | General function [2] | Osteoblast (OB) association | Osteoclast (OC) association |
|---|---|---|---|---|---|---|---|
| PRTN3 | - | + | n.s | Unclear | Serine protease. May play a role in antler shedding. | No direct evidence; may indirectly suppress osteoblast differentiation through PGRN degradation | No direct evidence; may indirectly facilitate osteoclastogenesis through loss of PGRN-mediated inhibition |
| ITIH6 | + | + | - | Unclear | Serine-type endopeptidase inhibitor activity | Putative promotion [3] | No direct evidence |
| COL8A1 | + | + | - | Unclear | Vastatin inhibits aortic endothelial cell proliferation and causes cell apoptosis | Putative promotion [4] | No direct evidence |
| COL16A1 | - | - | + | Unclear | Involved in mediating cell attachment and inducing integrin-mediated cellular reactions | Putative promotion (indirect evidence) [5] | No direct evidence |
| MSTN | + | + | - | OC | Acts specifically as a negative regulator of skeletal muscle growth | Inhibits osteoblast differentiation and mineralization [7] | Promotes osteoclastogenesis and osteoclast maturation [8] |
| ADPGK | - | n.s | + | Unclear | Catalyzes the phosphorylation of D-glucose to D-glucose 6-phosphate using ADP as the phosphate donor | Associated with osteoblast/osteocyte metabolic differentiation; no direct osteogenic evidence | No direct evidence |
| PON2 | - | n.s | + | Unclear | Capable of hydrolyzing lactones and a number of aromatic carboxylic acid esters. Has antioxidant activity | Indirectly supportive (antioxidant effect), but no direct evidence | Indirectly inhibitory (antioxidant effect), but no direct evidence |
| LAMB1 | - | n.s | + | OB | Binding to cells via a high affinity receptor | Promotes osteogenic differentiation through laminin–integrin signaling [9] | No direct evidence |
| LAMB4 | - | n.s | + | Unclear | Binding to cells via a high affinity receptor | Possible indirect association through laminin-family ECM signaling [10] | No direct evidence |
| ZYX | + | n.s | - | OB | Adhesion plaque protein. Binds alpha-actinin and the CRP protein | Promotes mechanosensitive osteoblast/osteocyte responses through focal adhesion signaling and cytoskeletal remodeling [11] | No direct evidence. |
| FRZB | + | n.s | - | Unclear | Wnt-protein binding. Regulate cell growth and differentiation in specific cell types | Context-dependent; can promote osteogenesis in some settings through Wnt modulation (linked to netrin-like domain) [12] | No direct evidence; indirect Wnt-mediated effects |
| APOA4 | - | n.s | + | Unclear | May have a role in chylomicrons and VLDL secretion and catabolism. Lipid binding | Possible indirect support of osteogenesis through anti-inflammatory and metabolic effects | No direct evidence |
| MATN4 | + | - | | Unclear | Major component of the extracellular matrix of cartilage | No direct evidence. Putative promotion. | No direct evidence |
| CNMD | + | - | | Unclear | Stimulate chondrocytes growth and inhibit the growth of vascular endothelial cells | Supports bone repair and endochondral ossification; no direct osteoblast evidence [13] | Associated with delayed osteoclastogenesis in CNMD-deficient fracture models [14] |
| SFRP5 | + | - | | Unclear | Wnt-protein binding. Regulate cell growth and differentiation in specific cell types | Inhibits osteogenic differentiation through Wnt/β-catenin suppression [15] | No direct evidence |
| CILP | + | - | | Unclear | Probably plays a role in cartilage scaffolding. Overexpression may lead to impaired chondrocyte growth and matrix repair and indirectly promote inorganic pyrophosphate (PPi) supersaturation in aging and osteoarthritis cartilage. | No direct evidence; possible indirect effects via IGF-1/TGF-β signaling | No direct evidence |
| BMP4 | + | - | | OB-OC | Involve in skeletal system development | Strong promoter of osteoblast differentiation and mineralization [16] | Associated with osteoclastogenesis and bone remodeling [17] |

| | | | | | | |
|---|---|---|---|---|---|---|
| TGFB3 | + | - | 🟪 OB-OC | Ossification involved in bone remodeling | Regulates osteoblast proliferation, differentiation, and ECM synthesis [18] | Regulates osteoclastogenesis and bone remodeling [19] |
| CCN2 | + | - | 🟪 OB-OC | Promotes proliferation and differentiation of chondrocytes | Promotes osteoblast proliferation, differentiation, and matrix production [20] | Promotes osteoclastogenesis via DC-STAMP and RANK/OPG signaling [21] |
| TNXB | + | - | ⬜ Unclear | Substrate-adhesion molecule that appears to inhibit cell migration | No direct evidence; osteoblast activity largely unaffected [22] | Inhibits osteoclast multinucleation and bone resorption [22] |
| KERA | + | - | 🟥 OB | May be important in developing and maintaining corneal transparency and for the structure of the stromal matrix | Promotes osteoblast differentiation, mineralization, and bone formation [23] | No direct evidence |
| MATN1 | + | - | ⬜ Unclear | Cartilage matrix protein is a major component of the extracellular matrix of non-articular cartilage | No direct evidence; indirect role in skeletal development through cartilage ECM and growth plate function [24] | No direct evidence |
| LAMA3 | + | - | ⬜ Unclear | Differentiation of keratinocytes | Possible indirect osteogenic support through laminin–integrin signaling [25] | No direct evidence |
| TIMP2 | + | - | 🟥 OB | Complexes with metalloproteinases and irreversibly inactivates them by binding to their catalytic zinc cofactor | Promotes osteoblast differentiation and matrix mineralization [26] | Inhibits osteoclast-mediated matrix degradation and bone resorption [27] |
| TFPI2 | + | - | ⬜ Unclear | May play a role in the regulation of plasmin-mediated matrix remodeling | No direct evidence | Inhibits proteolytic ECM degradation associated with osteoclast-mediated resorption [28] |
| FST | + | - | 🟥 OB | Binds directly to activin and functions as an activin antagonist | Promotes osteoblast differentiation and bone formation via activin/myostatin inhibition [29] | Indirectly inhibits osteoclastogenesis through activin antagonism [30] |
| CHAD | + | - | 🟥 OB | Promotes attachment of chondrocytes, fibroblasts, and osteoblasts. May play an important role in the regulation of chondrocyte growth and proliferation | Promotes osteoblast adhesion, differentiation, and mineralization via α2β1 integrin signaling [31] | Inhibits preosteoclast motility and bone resorption via its integrin-binding domain [32] |
| CCL25 | + | - | 🟦 OC | Potentially involved in T-cell development | No direct evidence; may recruit mesenchymal stromal cells involved in skeletal repair. [33] | Promotes osteoclast formation and inflammatory bone resorption through CCR9-dependent p38/ERK signaling [34] |

MC: Metacarpal; n.s.: not significant; OB: Osteoblast; OC: Osteoclast; 🟥 osteoblast-promoting; 🟦 osteoclast-promoting; 🟪 both osteoblast- and osteoclast-associated; ⬜ unclear or insufficiently characterized dominant function

### Supplementary Datasets

**Supplementary datasets 1.** Material characterization data used for structural, compositional, and histological analyses.

**Supplementary datasets 2.** Normalized dataset. The dataset includes the protein abundance after normalization, and one-tailed t-test p-values and their combinations.
